# TFIIH kinase CDK7 antagonizes phenotype switching and emergence of drug tolerance in melanoma

**DOI:** 10.1101/2020.09.24.311431

**Authors:** Pietro Berico, Max Cigrang, Cathy Braun, Guillaume Davidson, Jeremy Sandoz, Stephanie Legras, François Peyresaubes, Carlos Mario Gene Robles, Jean-Marc Egly, Emmanuel Compe, Irwin Davidson, Frederic Coin

## Abstract

Melanoma cells switch back-and-forth between phenotypes of proliferation and invasion in response to changing microenvironment, driving metastatic progression. We show that inhibition of the TFIIH kinase CDK7 (CDK7i) results in a melanocytic to mesenchymal phenotype switching and acquisition of targeted therapy tolerance. We identify a gene expression program controlled by the transcription factor GATA6, which participates in drug tolerance in mesenchymal-like cells and which is antagonized by CDK7 in melanocytic-like cells. This program emerges concomitantly with loss of melanocyte lineage-specific MITF protein following CDK7i. By dissecting the underlying mechanism, we observe that CDK7 accumulates at the super-enhancer regulating MITF to drive its expression. MITF itself binds to a intronic region of GATA6 to transcriptionally repress it. This molecular cascade antagonizes expression of the GATA6 regulon that only emerges in MITF-low cells of metastatic melanoma. Our work reveals a role for CDK7 in counteracting phenotype switching and activation of a gene expression program mediating multidrug tolerance in melanoma cells.

## Introduction

Among the protein complexes essential for gene expression in eukaryotes, the basal transcription factor TFIIH is unique due to its various enzymatic activities, including helicase, translocase and kinase functions (Berico and Coin, 2018; Villicana et al., 2014). TFIIH comprises 7 core forming subunits (composed of the XPB translocase, the XPD helicase together with the p62, p52, p44, p34, and TTDA subunits) that associate with the CDK-Activating-Kinase (CAK) sub-complex (containing the CDK7 kinase together with MAT1 and Cyclin H) (Compe and Egly, 2016). The CDK7 kinase is a positive kinase that phosphorylates transcription factors to promote gene expression (Fisher, 2019). The main target of CDK7 is the carboxyl-terminal domain (CTD) of the largest subunit of RNA polymerase II (RNAPII) at serine 2 (ser-2) and 7 (ser-7) contributing to the proper progression of transcription initiation (Eick and Geyer, 2013; Fisher, 2019). Surprisingly, inhibition of the CDK7 kinase activity (CDK7i) has led to impressive responses in various cancers (Cao and Shilatifard, 2014; Christensen et al., 2014; Kwiatkowski et al., 2014) probably due to the participation of the TFIIH kinase to super-enhancer-linked oncogene transcription (Chipumuro et al., 2014).

Malignant melanoma is responsible for 70% of skin cancer deaths in western countries (Eggermont et al., 2014). The 5-year survival rate is greater than 90% for localized melanomas, but drops to 16% for distant-stage disease demonstrating that metastases are responsible for patient mortality (Balch et al., 2011). Somatic gain-of-function mutations in the proto-oncogene kinase *BRAF* are the commonest mutations (60%) and the T → A transversion underlying BRAF^V600E^ comprises the majority of *BRAF* mutations (Brose et al., 2002; Davies et al., 2002). As an alternative to *BRAF* mutations, human melanomas commonly carry *NRAS* and *NF1* mutations (35%), while a number of patients show no mutations of these three genes (Triple-wt) (5%) (Hodis et al., 2012).

Melanoma is notorious for its heterogeneity based on co-existing melanoma cell phenotypes. In cultured melanoma cells, gene expression analyses have established two main and distinct signatures defined as either proliferative (melanocytic-type) or invasive (mesenchymal-like) melanoma cell states (Carreira et al., 2006; Verfaillie et al., 2015; Widmer et al., 2012). At the transcriptional level, the differentiated melanocytic-type melanoma cells display high levels of lineage-specific transcription factors, including the *microphthalmia-associated transcription factor (MITF)* and *SRY-box 10 (SOX10),* while the undifferentiated mesenchymal-like melanoma cells show low levels of *MITF* and *SOX10* but high expression of markers such as the *AXL receptor tyrosine kinase (AXL)* and *SRY-Box 9 (SOX9).* The discovery of cells with intermediate signatures supports the initial concept of phenotypic plasticity, called phenotype switching, driving melanoma progression through conversion from one phenotype into another in response to external cues and which is reminiscent to epithelial-mesenchymal transition (EMT) (Ennen et al., 2017; Hoek et al., 2008; Rambow et al., 2018).

Treatment options for patients with metastatic melanoma have changed dramatically in the past 10 years. Combination therapies with inhibitors targeting BRAF (i.e Vemurafenib and Dabrafenib) and MEK (i.e Trametinib) kinases (BRAFi and MEKi, respectively) have emerged and show high efficiency but are limited by development of resistance and subsequent progression (Menzies and Long, 2014). Studies have shown that tolerance to targeted therapies can involve various phenotype changes including the EMT-like process of melanocytic to mesenchymal phenotype switching (Arozarena and Wellbrock, 2019; Kemper et al., 2014; Rambow et al., 2019; Rambow et al., 2018). Therefore, understanding the molecular detail of phenotypic plasticity of melanoma cells is crucial for the development of future therapeutic approaches.

Here we identify the CDK7 kinase activity of TFIIH as a repressor of phenotype switching in melanoma, antagonizing the emergence of transcription program involved in drug tolerance. By dissecting the link between CDK7, phenotype switching and drug tolerance, we identified a new transcriptional program in mesenchymal-like cells, under the control of the GATA-binding factor 6 protein (GATA6), which we show to be involved in multidrug resistance and to be antagonized by CDK7 activity in melanocytic-type melanoma cells. Mechanistically, we unveil a negative molecular cascade in which CDK7 drives the expression of MITF that binds to an intronic regulatory sequence of the GATA6 locus to transcriptionally repress it. Therefore, the loss of MITF during CDK7i and phenotype switching promotes the progressive activation of the GATA6-dependent transcription program. Consistently, we observe that the GATA6 regulon emerges in MITF-low mesenchymal-like melanoma cells of human metastatic melanoma.

## Results

### Melanoma cultures exhibit distinct sensitivity to CDK7i

To study the role of CDK7 in melanoma, we first explored the sensitivity of melanoma cells to the loss of CDK7 kinase activity by using unique patient-derived 2D cultures (MM011, MM029, MM047, MM074, MM099 and MM117) (Gembarska et al., 2012; Verfaillie et al., 2015) together with one long-term frequently used melanoma cell line (501mel). MM029 (BRAF^V600K^), MM047 (NRAS^Q61R^) and MM099 (BRAF^V600E^) showed a mesenchymal-like gene expression signature that included low levels of the lineage-specific transcription factor MITF and high levels of SOX9 and c-Jun (Figure 1a) (Verfaillie et al., 2015; Widmer et al., 2012). In contrast, the melanocytic-type melanoma cells 501mel (BRAF^V600E^), MM011 (NRAS^Q61K^), MM074 (BRAF^V600E^) and MM117 (Triple-wt) exhibited moderate to high expression of MITF and SOX10 together with undetectable levels of expression of SOX9 and c-Jun (Verfaillie et al., 2015; Widmer et al., 2012) (Figure 1a).

**Figure 1:**
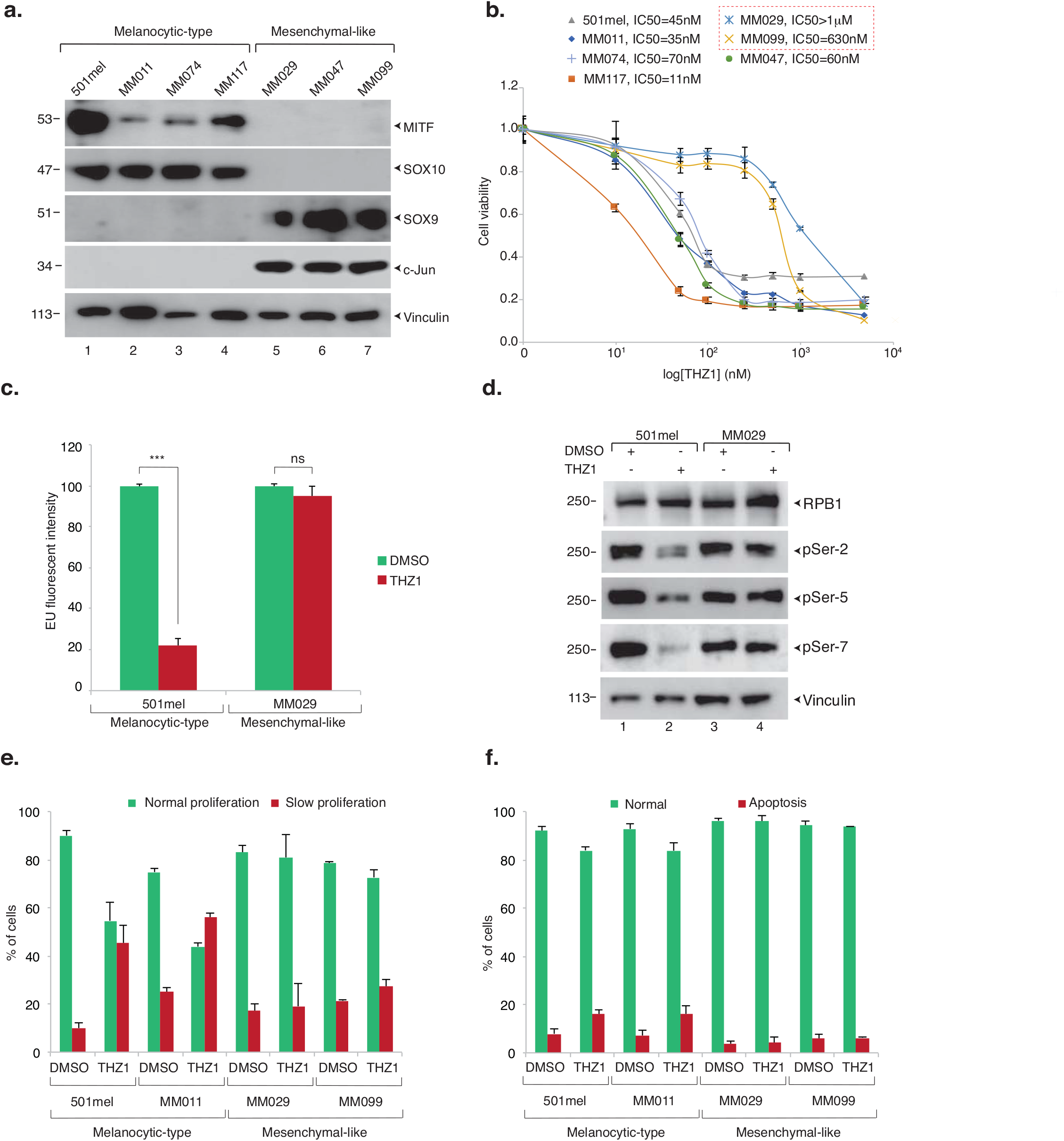
Mesenchymal-like melanoma cells resist to CDK7i. **a.** Protein lysates from either melanocytic-like (proliferative) 501mel, MM011, MM074 and MM117 or mesenchymal-like (invasive) MM029, MM047 and MM099 melanoma cell cultures were immuno-blotted for proteins as indicated. Molecular sizes of the proteins are indicated (kDa). **b.** Melanoma cell cultures were treated with increasing concentrations of THZ1 for 72 h. Mean growth is shown relative to vehicle (DMSO)-treated cells. Error bars indicate mean values + Standard Deviation (SD) for three biological triplicates. IC50 for each cell line is indicated. **c.** Melanocytic-type 501mel and mesenchymal-like MM029 melanoma cells were treated 4 h with either vehicle (DMSO, grey boxes) or THZ1 (50nM, black boxes). Transcribed RNAs were labelled by 5EU incorporation. Error bars indicate mean values + SD for three biological triplicates. N=~2500 cells were analyzed in each experiment (P-value ***<0.005, ns=non significant (>0.5), Student’s t-test). **d.** Protein lysates from melanocytic-type 501mel and mesenchymal-like MM029 melanoma cells, treated with vehicle (DMSO) or THZ1 (50nM) for 4 h, were immuno-blotted for proteins and for phosphorylation of ser-2, ser-5 and ser-7 of the CTD. Molecular sizes (KDa) of the proteins are indicated in kDa. **e-f.** Melanoma cells were treated with either vehicle (DMSO) or THZ1 (50nM) for 72 h. Cell proliferation and apoptosis were analysed using CellTrace **(e)** and Annexin V **(f)** staining respectively, followed by flow cytometry.

We next observed that the melanocytic-type 501mel, MM011, MM074, MM117 cells together with the mesenchymal-like MM047 cells were sensitive to low concentrations of the first-in-class CDK7 inhibitor THZ1 (Kwiatkowski et al., 2014) (Figure 1b) (IC50 ranging from 10 to 70nM). In marked contrast, the mesenchymal-like MM099 and MM029 cells were insensitive to CDK7i, even at high concentrations of the drug (IC50>500nM) (Figure 1b).

Because CDK7 plays a positive role in transcription initiation, we next measured the impact of CDK7i on global transcription by labeling *de novo* synthesized RNA with 5-ethynyl uridine (5EU) (Alekseev et al., 2017). While a global inhibition of RNA synthesis occurred in CDK7i-sensitive 501mel cells at low drug concentration, no transcriptional alteration was observed in CDK7i-insensitive MM029 cells after similar treatment (Figure 1c). Consistently, phosphorylation of initiation-associated ser-5 and ser-7 and elongation-associated ser-2 of the CTD of RNAPII was impaired in 501mel, while persisting in MM029 cells (Figure 1d). CDK7i was accompanied by an anti-proliferative effect on the 501mel and MM011 cells together with a slight induction of apoptosis, which was not observed in CDK7i-insensitive MM029 and MM099 melanoma cells (Figures 1e-f). Taken together, these observations demonstrated that melanocytic-type melanoma cells displayed high sensitivity to CDK7i, regardless of their driver mutation, while mesenchymal-like melanoma cells were mostly insensitive to it.

### CDK7 activity antagonizes melanoma phenotype switching

To unveil the molecular mechanisms controlling the sensitivity/insensitivity of melanoma cells to CDK7i, we generated THZ1 (CDK7i) or Vemurafenib (BRAFi) resistant cell lines *ex vivo.* To this end, we chronically exposed the melanocytic-type and CDK7i/BRAFi sensitive MM074(BRAF^V600E^) cells to escalating doses of the corresponding drug over several weeks. These treatments were carried out until the cells proliferated in drug concentrations equal to at least 5 times the IC50 values of the original cells, allowing us to generate the stable MM074^CDK7i-R^ and MM074^BRAFi-R^ cell lines, respectively (Supplemental Figures 1a-b). In parallel, the CDK7i-sensitive and mesenchymal-like MM047 cells (NRAS^Q61R^) were chronically exposed to THZ1 following the same protocol to generate stable MM047^CDK7i-R^ cells (Supplemental Figure 1c). Interestingly, establishment of CDK7i resistance decreased the sensitivity of the MM074^CDK7i-R^ cells to BRAFi (Vemurafenib) and MEKi (Trametinib) (Supplemental Figures 1b and d). In apparent contrast, the BRAFi-resistant MM074^BRAFi-R^ cells remained sensitive to both CDK7i and MEKi (Supplemental Figure 1a and d). Acquired resistance to CDK7i correlated with maintenance of global transcription synthesis of MM047^CDK7i-R^ and MM074^CDK7i-R^ cells upon CDK7i treatment when compared to their respective parental cells (Supplemental Figure 1e-f).

Using RNA-seq, we next analyzed the transcriptome of the CDK7i, BRAFi and parental cells. We first observed a pronounced modification of the transcriptional programs of MM074^CDK7i-R^ and MM074^BRAFi-R^ cells compared to MM074 cells and a less pronounced modification of the MM047^CDK7i-R^ cells compared to MM047 cells (Figure 2a). Analyses of the CDK7i-resistant cells showed deregulation of more than 6.000 genes in MM074^CDK7i-R^ cells compared to MM074 cells and 1.000 genes in MM047^CDK7i-R^ cells compared to MM047 cells (Figure 2b). Interestingly, despite the fact that the parental lines were of different cell types, 261 genes were commonly upregulated in the two CDK7i-resistant cell cultures (Figure 2b and Supplemental Data 1). As shown by Gene Ontology (GO) analysis, these genes were involved in epithelial cell differentiation or in the transport of small molecules, which could explain the resistance of melanoma cells to CDK7i (Supplemental Figure 2a).

**Figure 2:**
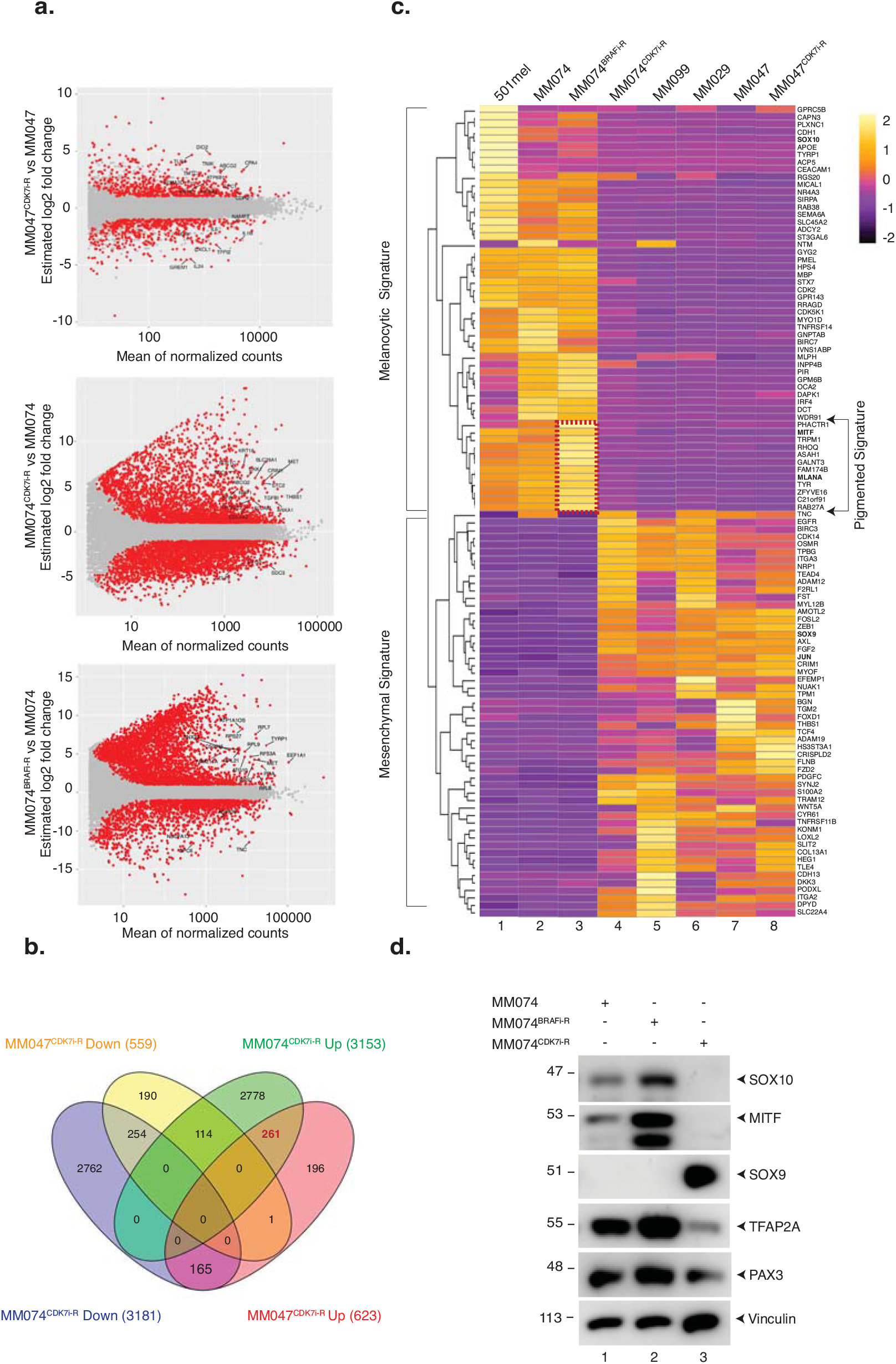
Exposure to CDK7i induces phenotype switching. **a.** Volcano plots were used to demonstrate differentially expressed genes as determined by RNA-seq in either MM047^CDK7i-R^ *vs* MM047 (top), MM074^CDK7i-R^ *vs* MM074 (middle) or MM074^BRAFi-R^ *vs* MM074 cells (bottom). Red dots show significantly over-(top) or under-(bottom) represented mRNA in drug-resistant cells compared to parental cells. All data were evaluated with the DESeq2 R package. The value for a given gene is the normalized gene expression value relative to the mean of all samples belonging to the same condition. **b**. Venn diagram indicating the number of up- and downregulated genes in MM047^CDK7i-R^ and MM074^CDK7i-R^ compared to the parental MM047 and MM074 cell lines. The number of genes overlapping between the four data sets is indicated. **c.** Genes characterizing the melanocytic-type and mesenchymal-like transcription signatures (Widmer et al., 2012) have been plotted in relation to their expression in different melanoma cell lines through a Heatmap where the RPKM values are represented as z-score. The group of genes related to pigmentation has been highlighted in red to show their over-expression in MM074^BRAFi-R^. Left, the color key showing the log2 expression values. Yellow color stands for high expression and dark violet for low expression. For reference, several genes including MITF, SOX10, MLANA, and SOX9, JUN are highlighted in blue. **d.** Protein lysates either from MM074, MM074^BRAFi-R^ or MM074^CDK7i-R^ were immuno-blotted for indicated proteins. Molecular sizes of the proteins are indicated (kDa).

We next conducted a bioinformatics approach to cluster the cell lines based on the expression of a hundred genes corresponding to previously described signatures of melanocytic *vs.* mesenchymal transcriptional cell states (Widmer et al., 2012). With this approach and in agreement with the literature (Verfaillie et al., 2015), the 501mel and MM074 cells showed a melanocytic-type transcriptional signature (Figure 2c, lanes 1-2), while the MM047, MM099 and MM029 cells showed a mesenchymal-like signature (lanes 5-7). Interestingly, we observed a mesenchymal-like signature in MM074^CDK7i-R^ cells, which differed from the melanocytic-type signature of the parental cells (compare lanes 2 and 4) and which correlated with an increased invasion capacity (Supplemental Figure 2b). In apparent contrast with MM074^CDK7i-R^, the melanocytic-type signature of the parental MM074 persisted in MM074^BRAFi-R^ cells in which we observed a significant increase in the expression of a set of *bona fide* pigmentation genes (Figure 2c, compare lanes 2 and 3), including *MLANA* (Supplemental Figure 2c), which correlated with higher cellular pigmentation (Supplemental Figure 2d).

The transcriptomic data prompted us to compare the amount of several protein markers of the melanocytic-type and mesenchymal-like states in the MM074, MMO74^CDK7i-R^ and MM074^BRAFi-R^ cells. We observed that the MM074^BRAFi-R^ cells exhibited higher amounts of the melanocytic lineage-specific proteins MITF, SOX10, TFAP2A and PAX3 compared to MM074 cells (Figure 2d). In contrast, MM074^CDK7i-R^ cells showed a dramatic loss of these proteins together with the emergence of SOX9 (Figure 2d).

In summary, melanoma cells chronically exposed to CDK7i adopted (for MM74^CDK7i-R^) or even heightened (for MM47^CDK7i-R^) an undifferentiated mesenchymal-like cell state, while cells exposed to BRAFi acquired a highly pigmented hyper-differentiated cell state.

### The GATA6 regulon is expressed in mesenchymal-like melanoma cells

Next, we analyzed the 261 common up-regulated genes in CDK7i-resistant cells (Supplemental Data 1) to identify a gene expression signature reflecting the observed CDK7i resistance. We first merged these genes with a list of annotated transcription factors and identified 16 common up-regulated transcription factors (TFs) in MM074^CDK7i-R^ and MM047^CDK7i-R^ cells (Supplemental Figure 3a). We next analyzed the expression of these TFs in melanoma cells *in vivo* using data from scRNA-seq performed on 674 cells derived from one drug-naive PDX tumor (Rambow et al., 2018). Seven out of the 16 TFs were expressed in a significant fraction of cells from the PDX (Supplemental Figure 3b). However, only the transcription factor *GATA6* was significantly overexpressed in primary melanoma compared to nevi (Supplemental Figure 3c). In line with these data, we observed high levels of *GATA6* mRNA and GATA6 protein in MM074^CDK7i-R^ but also in the CDK7i-insensitive MM029 and MM099 cells, compared to CDK7i-sensitive 501mel, MM011, MM117, MM074 cells (Figures 3a-b). We noticed low level of GATA6 protein in MM047, which however slightly increased in MM047^CDK7i-R^ (Figure 3b).

**Figure 3:**
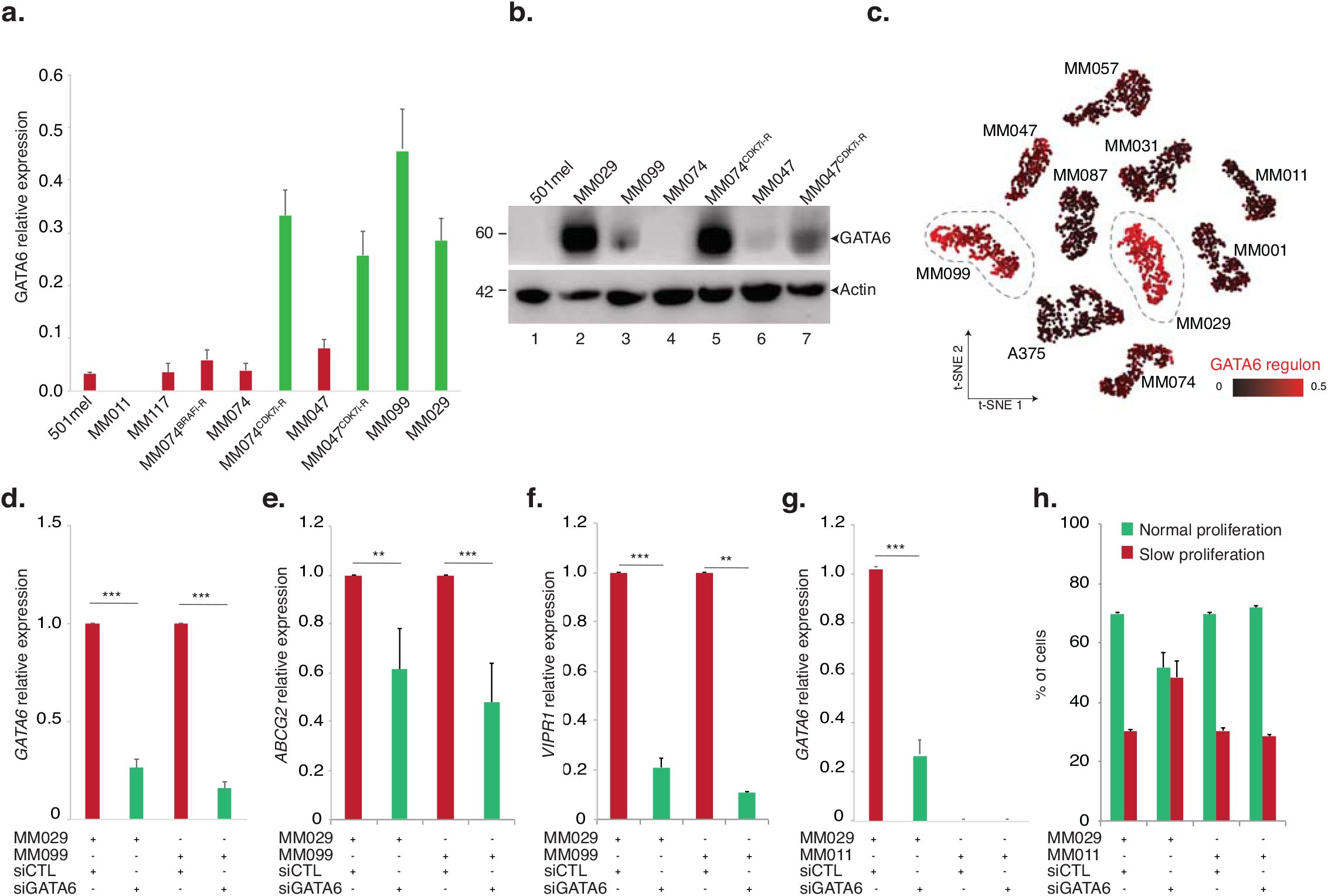
Differential expression of GATA6 regulon in melanoma cells. **a.** qRT-PCR analysis showing average *TBP*-normalized expression of *GATA6* in the indicated cell lines. Cells expressing GATA6 correspond to CDK7i-resistant cells (in green). Error bars indicate mean values + SD for three biological triplicates. **b.** Protein lysates from the indicated cell lines were immuno-blotted for the indicated proteins. Molecular sizes of the proteins are indicated in kDa. **c.** Seurat T-distributed stochastic neighbor embedding (t-SNE1/2) plots of the indicated melanoma cells colored based on the expressional activity of genes forming the *GATA6* regulon. Expression was retrieved from scRNA-seq of the melanoma cell cultures (GSE134432) (Wouters et al., 2020) and visualized with SCOPE. MM01 (Triple-wt), MM011, MM031 (BRAF^V600E^), MM057 (NRAS^Q61L^), MM074, MM087 (Triple-wt) and A-375 (BRAF^V600E^) are melanocytic-type melanoma cells. MM099, MM029 and MM047 are mesenchymal-like melanoma cells. MM099 and MM029 are marked by dashed ellipses. **d-f.** qRT-PCR analysis showing average *TBP*-normalized expression of *GATA6* **(d)***, ABCG2* **(e)** and *VIPR1* **(f)** in the indicated cell lines treated with either siCTL or si*GATA6* for 72 h. Error bars indicate mean values + SD for three biological triplicates. **g-h.** Melanocytic-type MM011 and mesenchymal-like MM029 melanoma cultures were treated with either siCTL or si*GATA6* for 72 h. qRT-PCR analysis shows average *TBP*-normalized expression of *GATA6* in the indicated cell lines **(g)**. Cell proliferation was analysed using CellTrace staining and flow cytometry in the indicated cell lines **(h)**. Error bars indicate mean values + SD for three biological triplicates.

We next used pySCENIC (Single-Cell rEgulatory Network Inference and Clustering) through SCOPE (scope.aertslab.org) on genome-wide transcriptome data recently obtained by scRNA-seq from shorted-cultured cells (Wouters et al., 2020). We observed that the GATA6 regulon, containing 359 genes (Supplemental Data 2), was highly enriched in the mesenchymal-like MM099 and MM029 cells (Figure 3c). Strikingly, depletion of GATA6 using short-interference RNA (siRNA) in MM029 and MM099 cells decreased expression of genes that were defined as part of the GATA6 regulon by pySCENIC, such as the efflux pump *ATP Binding Cassette Subfamily G member 2 (ABCG2)* and the *Vasoactive Intestinal Peptide Receptor 1 (VIPR1)* (Figures 3d-f). Furthermore, MM029 proliferation was significantly impacted upon GATA6 silencing compared to melanocytic-type MM011 that did not express GATA6 (Figures 3g-h).

### The efflux pump ABCG2 is involved in tolerance to CDK7i and BRAFi

We then compared the 261 common up-regulated genes in MM047^CDK7i-R^ and MM074^CDK7i-R^ with the GATA6 regulon and found 35 genes in common between the two signatures, amongst which was the efflux pump *ABCG2* (Supplemental data 1 and Supplemental Figure 4a). *ABCG2* belongs to the family of ATP-binding cassette (ABC) transporters playing a role in multidrug resistance (Robey et al., 2018). Analysis of RNA-seq data from melanoma tumors or *in situ* mRNA hybridization of melanoma tumor sections demonstrated higher expression *of ABCG2* in cutaneous metastatic melanoma compared to primary tumors (Supplemental Figures 4b-c). In addition, three ABC transporters *(ABCB1, ABCC3* and *ABCG2)* were up-regulated in MM047^CDK7i-R^ and/or MM074^CDK7i-R^ cells (Supplemental Figures 4d-e) but only *ABCG2* was significantly over-expressed in mesenchymal-like CDK7i-resistant MM099 and MM029 cells (Figures 4a-b). Depletion of ABCG2 using siRNA (Supplemental Figure 4f) significantly sensitized MM099 and MM029 to CDK7i (Figures 4c-d). Interestingly, depletion of ABCG2 also significantly sensitized the MM099 cells to BRAFi (Figure 4e), showing the pleiotropic impact of this efflux pump on drug resistance in these cells. Rationally, MM029 cells bearing the Vemurafenib-resistance associated BRAF^V600K^ mutation were insensitive to BRAFi with or without ABCG2 (Figure 4f). Taken together, these data suggested that the ABC transporter ABCG2 played a significant role in the tolerance to CDK7i and BRAFi in melanoma cells.

**Figure 4:**
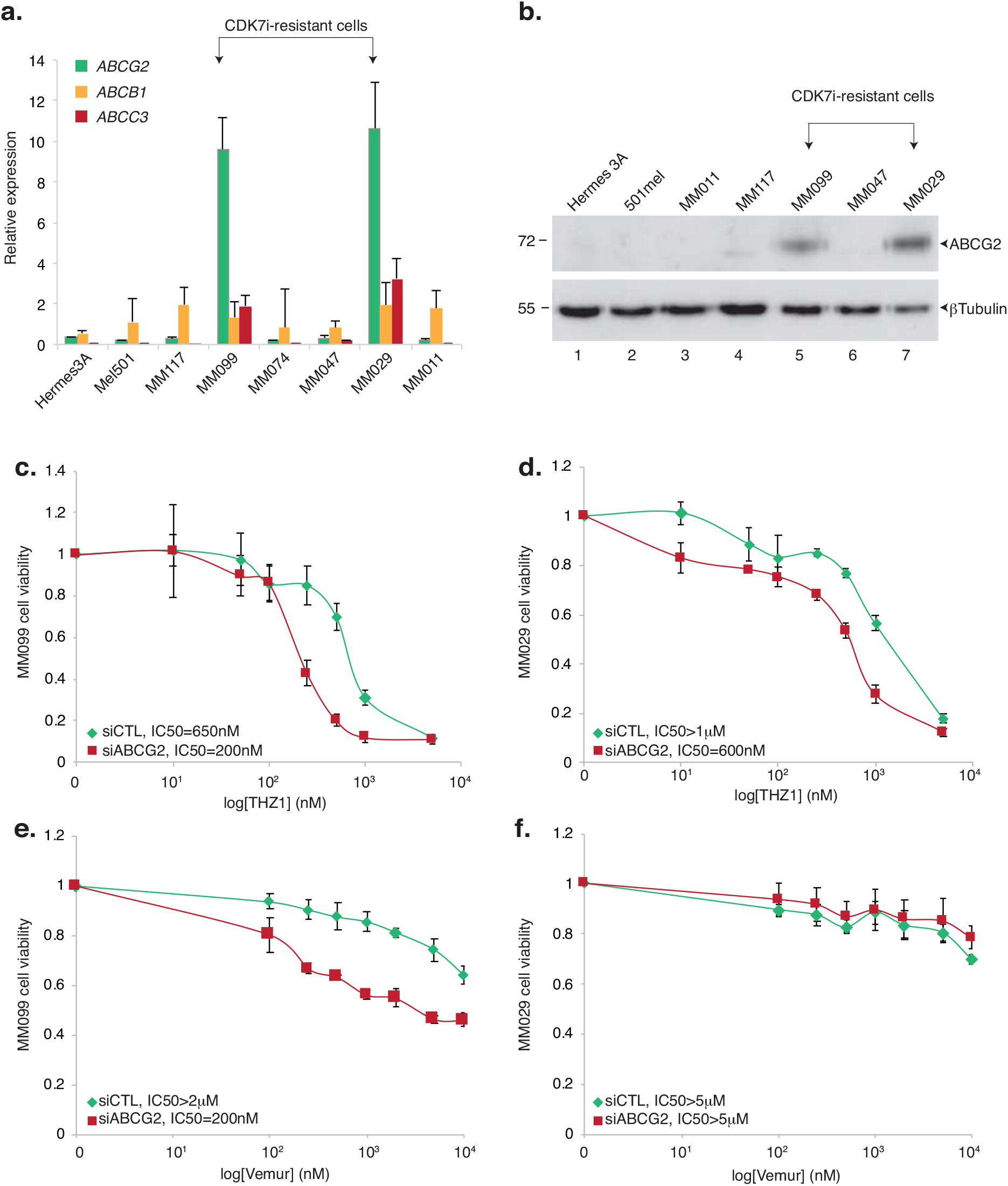
ABCG2 is involved in multidrug tolerance in melanoma cells. **a.** qRT-PCR analysis showing average *TBP*-normalized expression of *ABCB1, ABCC3* and *ABCG2* in the indicated cell lines. Error bars indicate mean values + SD for three biological triplicates. **b.** Protein lysates from the indicated cell lines were immuno-blotted for the indicated proteins. Molecular sizes of the proteins are indicated in kDa. **c-d.** MM029 **(c)** and MM099 **(d)** melanoma cells were pre-treated with either siCTL or *siABCG2* as indicated and treated with increasing concentrations of THZ1 for 72 h. Mean growth is shown relative to vehicle (DMSO)-treated cells. Error bars indicate mean values + SD for three biological triplicates. IC50 for each cell line is indicated. **e-f**. MM029 **(e)** and MM099 **(f)** melanoma cells were pre-treated with either siCTL or si*ABCG2* as indicated and treated with increasing doses of Vemurafenib for 72 h. Mean growth is shown relative to vehicle (DMSO)-treated cells. Error bars indicate mean values + SD for three biological triplicates. IC50 for each cell line is indicated.

### Loss of MITF induces expression of GATA6

We next sought to understand how CDK7 activity could counteract the emergence of the GATA6 regulon in melanocytic-like melanoma cells. Previous work suggested that CDK7 may occupy SE regulating MITF and SOX10 expression in melanoma cells {Eliades, 2018 #8337}. Using 501mel in which the two alleles coding for CDK7 were tagged with a Biotin-3xFlag tag by genome editing (501mel^BIO-FLAG:CDK7^) (Supplemental Figures 5a-b), we first aimed at identifying the chromatin binding profile of CDK7 in the melanocyte-type melanoma cells. Following FLAG Chip-seq, we identified numerous CDK7-binding sites throughout the locus of the lineage-specific *MITF* oncogene and of its transcriptional activator *SOX10* (Figure 5a-b). Binding of CDK7 occurred together with marks of H3K27ac, binding of MITF and/or of SOX10, BRG1 or H2AZ, all characterizing super-enhancer-type elements. In agreement with a role of CDK7 in MITF-SOX10 expression, a short 24 h CDK7i treatment impaired the expression of *MITF* and *SOX10* in 501mel cells (Figure 5c). Interestingly, loss of *SOX10* and *MITF* occurred in parallel with an increased expression of *GATA6* (Figure 5c). To strengthen this observation we calculated the correlation of *GATA6* with *MITF* or CDK7 expression in published RNA-seq data from human skin cutaneous melanoma (TCGA) and identified a consistent anti-correlation (r=-0.250 for *GATA6 vs. MITF* and r=-0.181 for *GATA6 vs. CDK7)* between these genes (Figure 5d).

**Figure 5:**
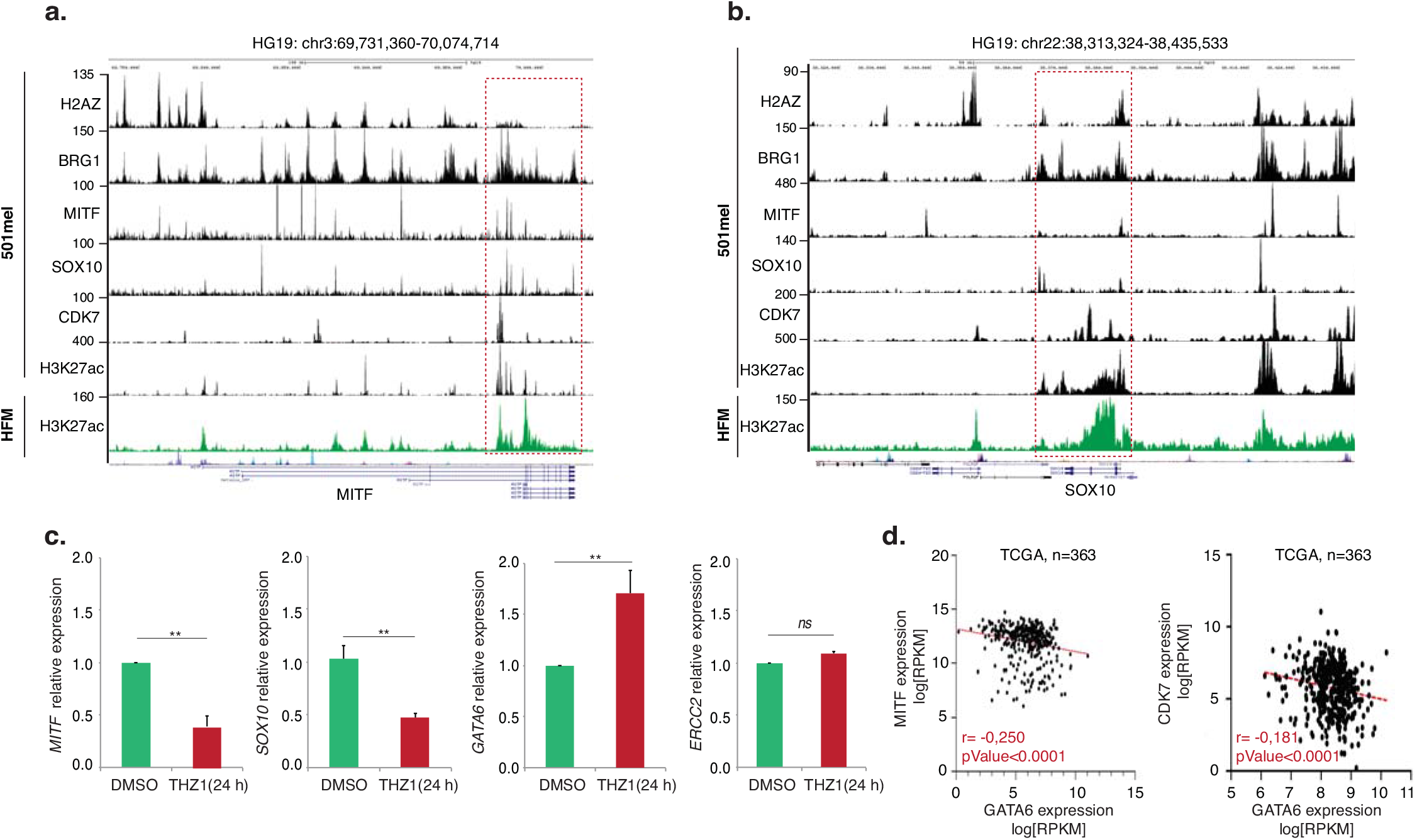
MITF and GATA6 expressions are anticorrelated. **a-b.** Gene track of CDK7 occupancy at *MITF* (**a**) and its transcriptional regulator *SOX10* (**b**) loci in 501mel^BIO-FLAG:CDK7^ cell line. Gene tracks of H2A.Z, BRG1, MITF, SOX10 and H3K27ac (GSE94488 and GSE61967) at the same loci in parental 501mel are indicated. SE is denoted by a red opened square. H3K27ac deposition is also shown in Hair Follicle Melanocytes (HFM) (GSE94488). **c.** qRT-PCR analysis showing average *TBP*-normalized fold expression of *SOX10, MITF, GATA6* and ERCC2 in melanocytic-type 501mel cells treated either with DMSO or THZ1 (50nM) for 24 h. Error bars indicate mean values + SD for three biological triplicates. p-value, **<0.05, ns=non significant (>0.5), Student’s t-test. Expression of *ERCC2* is used as a control unaffected by CDK7i treatment. **d.** Scatter plot expression values (in log[RPKM]) of *GATA6* against *MITF* (left) or *CDK7* (right) shows inverse relationship between GATA6 and these two genes in human skin cutaneous melanoma. These data were extracted from The Cancer Genome Atlas bulk RNA-seq data on a cohort of patient melanomas (n=363, Pan-Cancer Atlas). The line of best fit, the Spearman’s rank correlation coefficient (r) and the P-value (T-test) are indicated.

To assess if loss of MITF could mimic CDK7i and induces GATA6 expression, we directly depleted MITF or its transcriptional activator SOX10 with siRNA in 501mel cells and observed a significant induction of *GATA6* expression (Figure 6a). To analyse the expression of the whole GATA6 regulon after loss of MITF, we conducted a bioinformatic approach to exploit published scRNA-seq performed at different times after depletion of the MITF activator SOX10 in the CDK7i-sensitive MM074 cells (Wouters et al., 2020). We observed that the progressive down-regulation of the SOX10 regulon, containing MITF (Supplemental data 3), correlated with the progressive up-regulation of that of GATA6 (Figure 6b and Supplemental Figure 6a). We observed similar results with the melanocytic-type MM087 (Triple-wt) cells (Figure 6b and Supplemental Figure 6b). Altogether these data demonstrate an antagonism between GATA6 and MITF-CDK7 in melanoma both *ex vivo* and *in vivo*.

**Figure 6:**
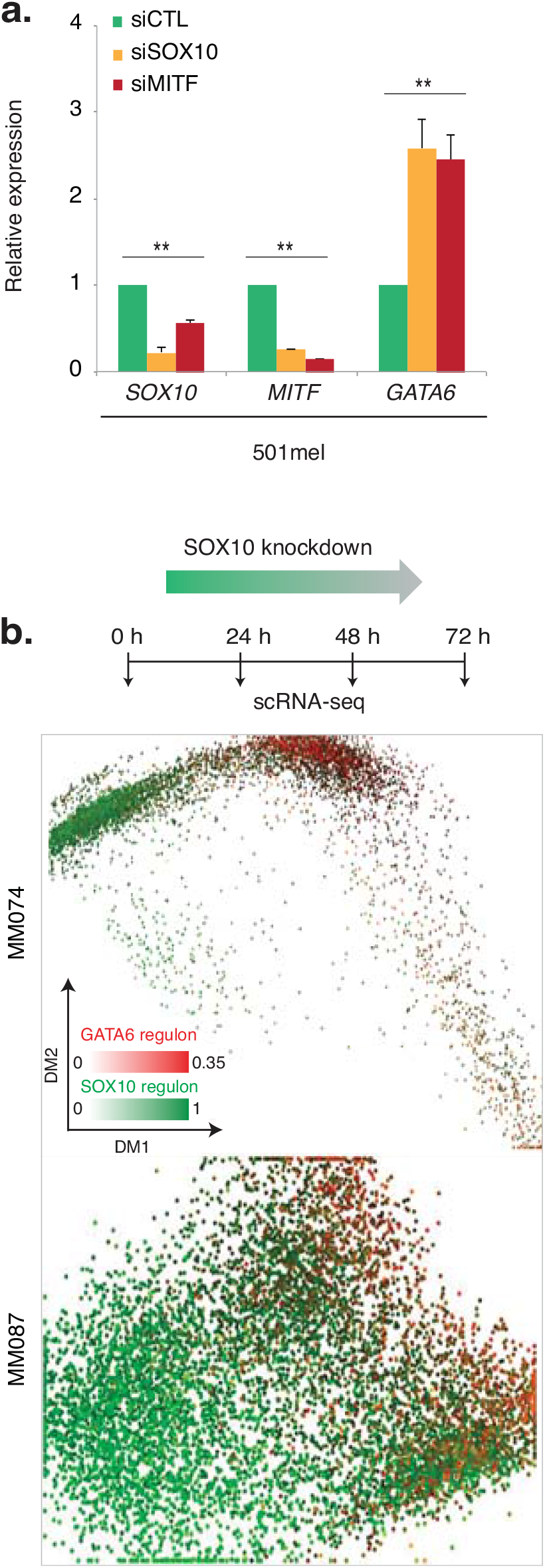
Loss of MITF releases *GATA6* expression. **a**. qRT-PCR analysis showing average *TBP*-normalized fold expression of *MITF, SOX10* and *GATA6* in melanocytic-type 501mel cells treated either with siCTL, *siSOX10* or *siMITF* for 48 h. Error bars indicate mean values + SD for three biological triplicates. p-value siCTL *vs* siSOX10 or siMITF, **<0.05, Student’s t-test. **b.** Diffusion map (DM1/2) with melanocytic-type MM074 (left panel) and MM087 cells (right panel) colored by expression of *SOX10-* (green) or *GATA6-* (red) regulons. The graphs show the trajectory that melanoma cells follow from 0 h (left) to 72h (right) post-si*SOX10* treatment as determined by scRNAseq at 0, 24, 48 and 72 h (GSE116237) (Wouters et al., 2020). An experimental setup is indicated on the left.

### MITF transcriptionally represses GATA6

The above observations prompted us to hypothesize a mechanistic link between MITF and the repression of *GATA6* in melanocytic-like cells. We analysed our published MITF ChIP-seq data set (Laurette et al., 2015) to determine potential MITF binding profile in the genomic region of *GATA6.* In an intronic region of the *GATA6* gene body, we identified a MITF-binding site containing also 3 E-box motifs that are, with M-box motifs, one of the two DNA recognition sequences of MITF (Figure 7a)(Laurette et al., 2015). This binding site was enriched in H3K27ac, BRG1 and H2AZ, marks of enhancer elements. Interestingly, in mesenchymal-like MM099 cells where MITF is not expressed, intronic H3K27ac was lost, but we observed a strong H3K27ac deposition at the *GATA6* promoter, which correlates with its high expression in these cells (Figure 7a). ChIP-qPCR of MITF and H3K27ac mark confirmed their enrichment in the E-box rich intronic region of the *GATA6* gene body in 501mel cells (Figure 7b).

**Figure 7:**
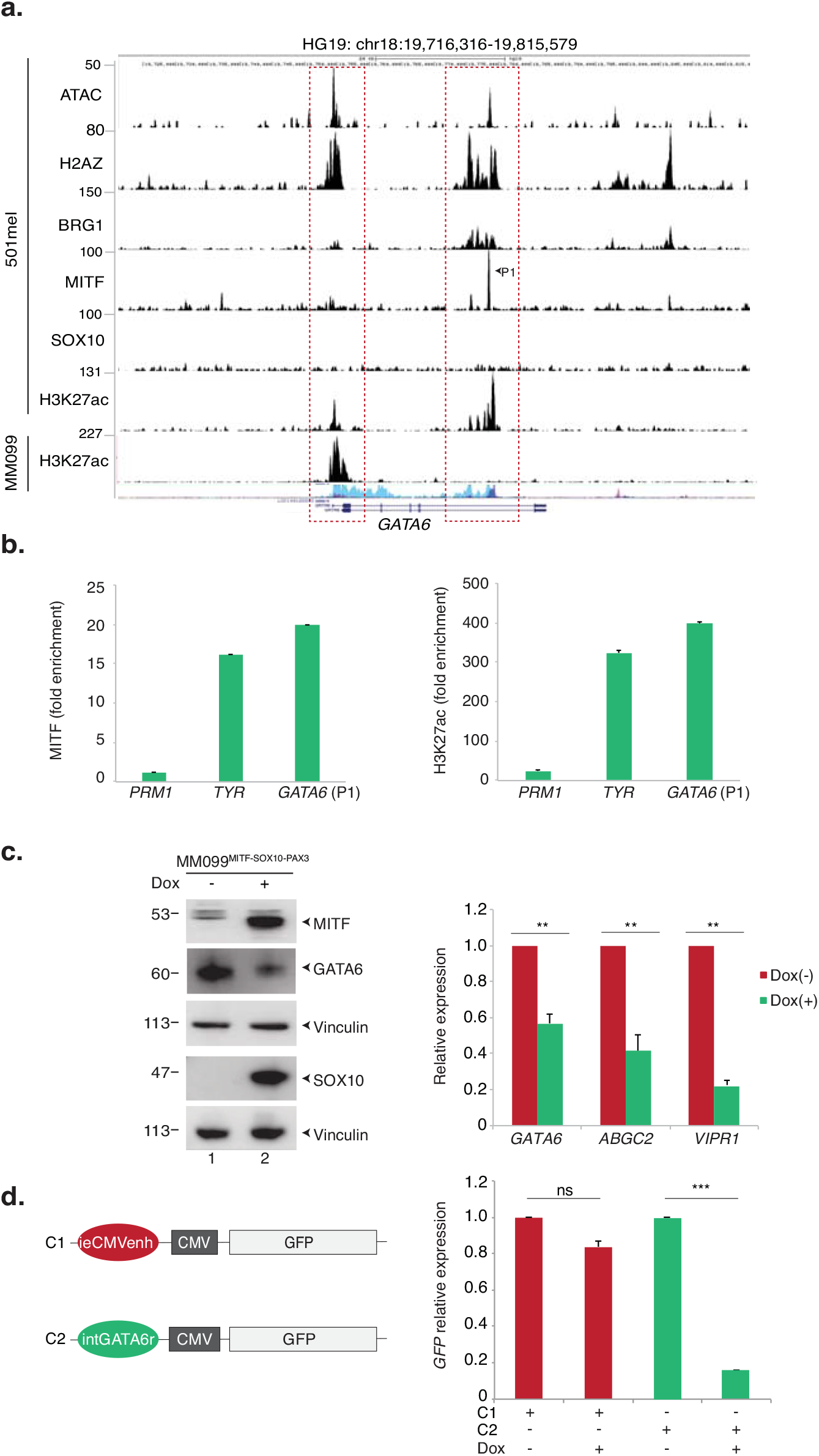
MITF binds GATA6 locus and transcriptionally represses it. **a.** Gene track of 3HA-MITF signal occupancy showing a significant MITF binding peak (P1) in the GATA6 gene body in the 501mel cell line (GSE61967). Additional tracks indicate potential regulatory regions highlighted by ATAC-seq and H3K27ac, BRG1, SOX10 and H2A.Z deposition (GSE94488 and GSE61967). H3K27ac deposition is also shown in mesenchymal-like MM099 cells at the *GATA6* locus. The scale bar indicates the size of the genomic region in kilobases (Kb). **b.** ChIP experiment monitoring the fold enrichment (compare to control IgG) of MITF protein and H3K27ac mark at the P1 *GATA6* intronic region identified above. *Proteamine 1 (PRM1)* and Tyrosinase *(TYR)* regulatory regions were used as negative and positive controlled, respectively (Laurette et al., 2015). **c. Left panel**; Mesenchymal-like MM099^MITF-SOX10-PAX3^ cells expressing inducible *MITF-SOX10-PAX3* genes were treated or not with Doxycycline (1μg/ml) for 24 h and protein lysates were immunoblotted for the indicated protein. **Right panel**; qRT-PCR analysis showing average *TBP*-normalized fold expression of *GATA6, ABCG2* and *VIPR1* in Mesenchymal-like MM099 cells treated or not with Doxycycline (1μg/ml) for 24 h. Error bars indicate mean values + SD for three biological triplicates. p-value **<0.05, ns=non significant (>0.5), Student’s t-test. **d. Left panel**; Schematic representation of pCDNA-ieCMVenh-CMV-GFP (C1) or pCDNA-intGATA6r-CMV-GFP (C2) reporter vectors. The ieCMVenh sequence in C1 was replaced by the intGATA6r sequence to generate C2. **Right panel**; qRT-PCR analysis showing average *TBP*-normalized fold expression of *GFP* in Mesenchymal-like MM099^MITF-SOX10-PAX3^ transfected with C1 or C2 vectors for 48 h before treatment or not with Doxycycline (1μg/ml) for 24 h. Error bars indicate mean values + SD for three technical triplicates. p-value ***<0.005, ns=non significant (>0.5), Student’s t-test.

To determine whether MITF was able to transcriptionally repress *GATA6,* we generated the MM099^MITF-SOX10-PAX3^ cell line in which MITF, SOX10 and PAX3 expression could be induced by doxycycline treatment (Figure 7c, left panel). Following induction of MITF-SOX10-PAX3, we observed a repression of *GATA6* mRNA level and a loss of GATA6 protein, which correlated with the repression of *ABCG2* and *VIPR1,* belonging to the GAAT6 regulon(Figure 7c).

In an attempt to demonstrate the repressive role of the regulatory intronic GATA6 region, the 500bp sequence encompassing the three E-box recognized by MITF (hereafter called “intGATA6r”) was subcloned upstream of the CMV promoter of the pcDNA-CMV vector to replace the immediate early CMV enhancer (ieCMVenh) in the context of a GFP reporter vector (Figure 7d, left panel). The reporter construct was transiently transfected into MM099^MITF-SOX10-PAX3^ expressing or not MITF-SOX10-PAX3. While the expression of the CMV under the ieCMVenh sequence was barely affected by the induction of MITF-SOX10-PAX3 expression, the presence of the intGATA6r element upstream the promoter decreased its activity ~ 85%, compared to cells that did not expressed MITF-SOX10-PAX3 (Figure 7d, right panel). Altogether, these data demonstrate that MITF transcriptionally represses *GATA6* by binding to a negative transcriptional regulatory sequence located in an intronic region of *GATA6*.

### GATA6 is enriched in undifferentiated melanoma cells *in vivo*

The above data imply that the expression of the GATA6 regulon correlates with MITF-low undifferentiated melanoma cells in tumors. To test this hypothesis, we first performed an immunohistological examination of human tumor samples and observed that GATA6 was highly expressed in a subpopulation of cells in cutaneous metastases, which did not express SOX10 (Figure 8a, panels e-f). In contrast, GATA6 was hardly observed in nevi and primary melanomas, which showed high expression of SOX10 (Figure 8a, panels a-d). Consistently, we observed higher expression of *GATA6* in metastatic melanoma compared with primary melanoma by analyzing public DNA microarray (Xu et al., 2008) or RNA-seq (TCGA) data (Supplemental Figure 7).

**Figure 8:**
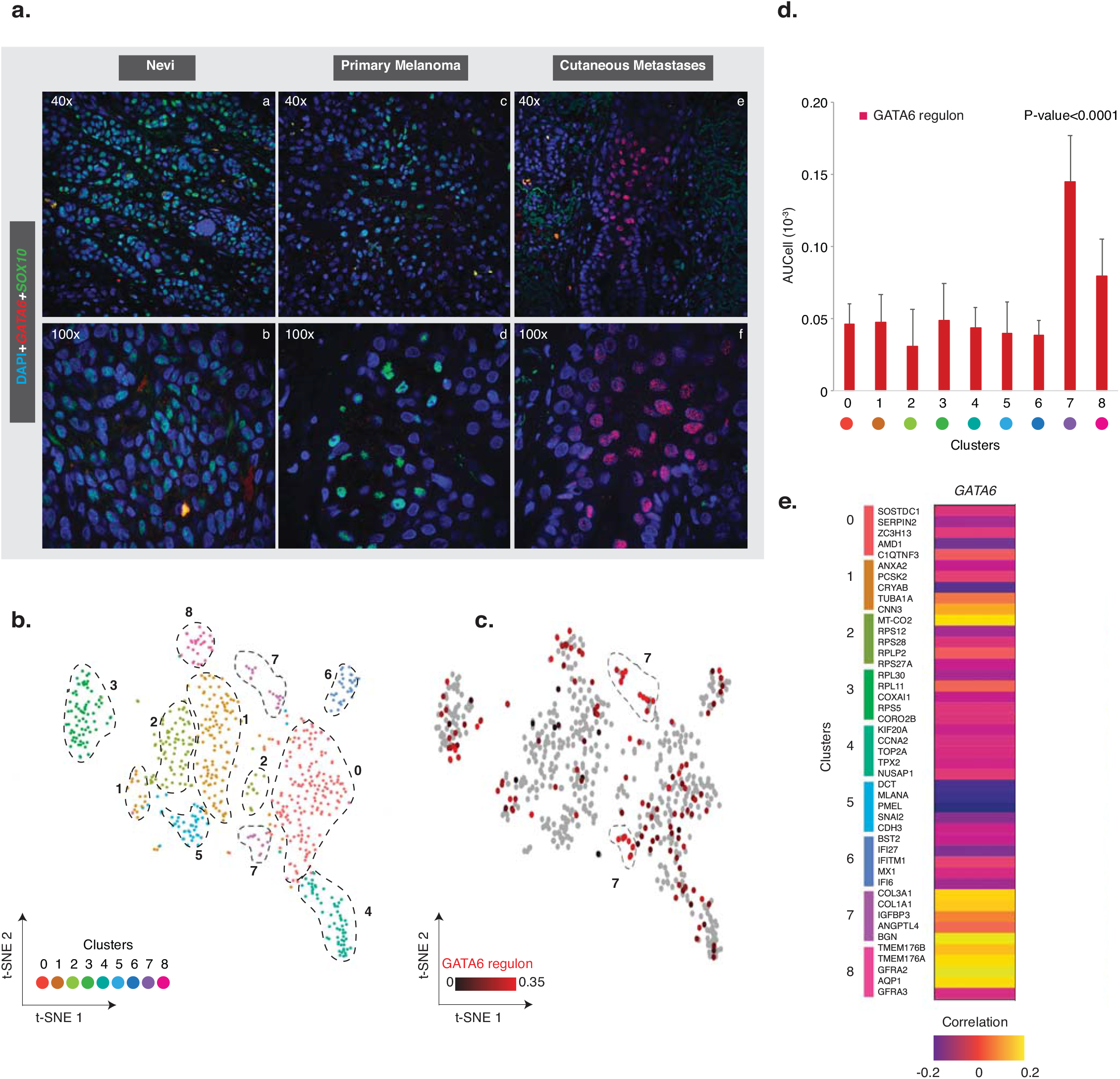
The GATA6 regulon is expressed in undifferentiated cells of melanoma tumours. **a.** Tumor sections were immuno-labelled with anti-GATA6 (red) and anti-SOX10 (green) antibodies and images were captured by confocal microscopy at the indicated magnification. **b.** Seurat t-SNE1/2 with cells colored according to their cluster. Data was retrieved from scRNA-seq performed on 674 cells from a single drug-naïve PDX (GSE116237) (Rambow et al., 2018) and visualized with SCOPE. **c.** Seurat t-SNE1/2 with cells colored in red scale according to the level of expression of the gene set forming the GATA6 regulon. Expression was measured by AUCell from scRNA-seq of PDX data used in (b) and visualized with SCOPE. **d.** The mean expression of the gene set forming the GATA6 regulon for each cluster is represented. Expression was measured by AUCell from scRNA-seq of PDX data used in (b) and reported on the graph. Error bars indicate mean values + SD. The P-value (One-way ANOVA) of cluster 7 *vs*. all is indicated. **e.** Heatmap illustrating the Spearman’s rank correlation coefficient between expression of *GATA6* and that of the indicated cluster markers extracted from TCGA RNA-seq of a cohort of patient melanomas (skin cutaneous melanoma, n=444) (Pan-Cancer Atlas) (Supplemental data 4). Yellow color stands for high correlation and dark violet for low correlation.

To further decipher which subtypes of melanoma cells may express *GATA6* and its regulon in a tumor, we used the Seurat script to conduct an unsupervised gene clustering analysis of published scRNA-seq data of the drug-naive PDX tumour mentioned above (Rambow et al., 2018). Our analyses allowed us to separate the 674 single cells in 9 different clusters (Figure 8b and Supplemental Figures 8a-b). According to the GO analysis performed for each cluster, a significant percentage of *GATA6* positive cells in the PDX belonged to cluster 7, which was characterized by genes involved in cell migration or RTK signalling pathways expressed in undifferentiated mesenchymal-like melanoma cells (Supplemental Figure 8b-c). In agreement, expression of the GATA6 regulon merged significantly with cluster 7 of mesenchymal-like melanoma cells (Figures 8c-d). Finally, using the TCGA RNA-seq cohort of patient melanoma mentioned above, we observed a positive correlation between the expression of *GATA6* and specific mesenchymal-like (cluster 7) and neural-crest cell (cluster 8) markers, expressed in undifferentiated melanoma cells (Figure 8e and Supplemental data 4). Altogether, these data establish that, as predicted from our *ex vivo* data, *GATA6* and its regulon are significantly enriched in MITF-low undifferentiated melanoma cells *in vivo*, such as the mesenchymal-like or neural-crest cells.

## Discussion

Here our report provides new insights into understanding the processes that govern gene regulatory network expression in melanoma by showing that a negative molecular cascade involving CDK7, MITF and GATA6 promotes the repression of a transcription program involved in drug tolerance in melanoma. To demonstrate this, we first studied CDK7i sensitivity of melanoma cells using patient-derived 2D-cultures that represent a better model system than classical long-term melanoma cell lines. This approach allowed us to observe variability in the CDK7i sensitivity of melanoma cells, which is independent of driver mutation status, but rather reflects their phenotype. Melanocytic-type melanoma cells are highly sensitive to CDK7i, which potently prevents their growth at low concentrations of THZ1. Contrary to melanocytic-type cells, mesenchymal-like melanoma cells were largely insensitive to CDK7i. The establishment of acquired CDK7i-resistant melanoma cells further allowed us to define a set of 260 genes that are commonly up-regulated upon CDK7i exposure. The altered expression of these genes, involved in epithelial cell and skin differentiation or in the transport of small molecules, reflects an adaptation to CDK7i treatment. Functional genomics allowed us to show that, among these genes, CDK7i provokes the emergence of a new transcription program in melanoma cells controlled by the transcription factor GATA6. We observed that the emergence of the GATA6 regulon following CDK7i exposure is instigated by decommission of the super-enhancers regulating the expression of the lineage-specific factor *MITF* and its regulator *SOX10* (Eliades et al., 2018). Indeed, we observed accumulation of CDK7 at these super-enhancers and we were able to mimic the effect of CDK7i on the emergence of the GATA6 regulon by depleting individually SOX10 or MITF proteins. Reciprocally, ectopic expression of MITF in mesenchymal-like cells led to the inhibition of *GATA6* expression, indicating a direct impact of MITF on *GATA6* expression. Ultimately, we unveiled the molecular detail of this inhibition cascade. Indeed, we observed that MITF binds to a short regulatory sequence located in an intron of the GATA6 locus. When introduced upstream of a reporter gene, the regulatory sequence repressed its transcription only in the presence of MITF. Then, and although the regulatory element displays the characteristics of an enhancer, it rather appears to be a negative regulatory element of GATA6 expression in melanoma cells when it is bound by MITF (Riesenberg et al., 2015). Of note, this is the first cloning of a *bona fide* negative regulatory sequence involving MITF. How might MITF exert a negative regulatory function by binding to the intGATA6r element remains to be determined but it seems likely that repressors interact with MITF in these conditions to turn-off the expression of *GATA6.* In any cases, our work reveals a molecular cascade of inhibition involving CDK7, MITF and GATA6 validated by the anti-correlation observed between MITF and GATA6 but also between CDK7 and GATA6 in tumors. Since MITF has been shown to participate to the stabilization of CDK7 in melanocyte-type melanoma cells (Seoane et al., 2019), our data suggest that the progressive loss of MITF during phenotype switching *in vivo* provokes the loss of CDK7 protein, which in turn will amplify the loss of MITF to result in the release of GATA6 expression in MITF/CDK7-low mesenchymal melanoma cells.

GATA6 is one of six members of the mammalian GATA family of transcriptional regulators and is expressed in various normal tissues derived from the mesoderm and endoderm (Almalki and Agrawal, 2016). An oncogenic role for GATA6 has been proposed in various cancers including pancreatic cancer where its knockdown reduced cell proliferation and cell-cycle progression (Sun and Yan, 2020). In line with these observations, we show that the loss of GATA6 impairs proliferation of mesenchymal-like cells. Since *GATA6* is expressed in normal adult tissues, it is unlikely that its targeting would lead to efficient therapy. However, this new transcription program may help to identify important molecular targets in mesenchymal-like melanoma cells, which could be exploited therapeutically to prevent acquisition of metastatic and drug resistance potential. Among these targets, we observed that the multidrug transporter *ABCG2* is responsible for cross-resistance to targeted therapies in mesenchymal-like cells. Because we observed that the GATA6 regulon, including *ABCG2, is* significantly overexpressed in metastatic melanoma tumors compared with primary tumors, these genes may mediate ubiquitous cross-resistance to targeted therapies clinically.

Together with the de-repression of gene expression programs, our results clearly established that melanocytic-type melanoma cells exposed to CDK7i progressively switch from a melanocytic to a mesenchymal-like cell state, suggesting a role for CDK7 in cell plasticity. It has been suggested that cellular phenotype changes, such as cell morphology or invasion capacity, leading to an invasive behavior of melanoma cells are rapid and appear before full establishment of the mesenchymal-like signature at the transcriptional level. For instance, the over-expression of the mesenchymal-like marker *SOX9* induces increased invasion *in vitro,* but this is only accompanied by a partial enrichment of invasive signature genes (Cheng et al., 2015). In acquired CDK7i-resistant melanoma cells, we detected an established mesenchymal-like transcriptional signature with, for instance, the acquisition of programs responsible for invasion. These observations show that these cells undergo a complete and stable phenotype switching characterized by a transcriptional reprogramming reminiscent of EMT.

In apparent contrast, we observed that the acquired resistance of melanocytic melanoma cells to BRAFi was not accompanied by a loss of lineage-specific transcription factors. Different outcomes of chronic exposure of melanoma cells to BRAFi exist in the literature (Rambow et al., 2019), depending probably on the starting cells but also on the strategy that is used to obtain the resistant cells (escalating doses *vs.* a single killing dose). In our hands, and as previously observed *in vitro* as well as in tumors (Smith et al., 2016)^-^(Haq et al., 2013), the chronic exposure of melanocytic-type melanoma cells to escalating doses of BRAFi switched them to a highly pigmented state characterized by an increased expression of pigmentation pathway-associated genes. Such hyper-differentiation is a likely consequence of the increased MITF expression that we observed in these cells (Khaled et al., 2010). Nevertheless, our data indicate that similar chronic exposure of a unique melanocytic-type melanoma cell culture to escalating doses of CDK7i or BRAFi promotes opposite effects with hyper-differentiation for BRAFi and undifferentiation for CDK7i.

Finally, many studies identified CDK7 as a cancer therapeutic target (Fisher, 2019; Sava et al., 2020). However, the phenotype switching observed during prolonged exposure of melanoma cells to CDK7i illustrates the potential danger of targeting this kinase in cancers in which EMT exists, a point that has never been investigated so far. Therefore, further studies should take into consideration the potential emergence, during CDK7i exposure of tumor cells, of mesenchymal-like cells able to resist to conventional therapies.

## Supporting information

Supplemental informations

## Acknowledgments

We thank the IGBMC antibody and cell culture facilities, Prof D. Lipsker and the staff of the Strasbourg University Hospital dermatology clinic for tumour sections and Dr Ghanem Ghanem for providing us the MM0-series melanoma cultures. This study was supported by the INCA (2017-11537), the Ligue contre le Cancer (Equipe labélisée 2019, FC and équipe labélisée 2018, ID) and the ANR-10-LABX-0030-INRT, a French State fund managed by the Agence Nationale de la Recherche under the frame program Investissements d’Avenir ANR-10-IDEX-0002-02. Sequencing was performed by the IGBMC GenomEast platform, a member of the “France Génomique” consortium (ANR-10-INBS-0009). P.B is supported by the Ligue contre le Cancer. M.C is supported by the Fondation pour la Recherche Médicale.

## Author Contributions

P.B., I.D and F.C. conceived the study. P.B., I.D, F.C and E.C analyzed the data. P.B. generated the resistant cells, performed the RNA-seq and the IH staining. S.L., G.D. and P.B., performed the bioinformatics analyses. P.B and C.M.G.R performed the survival assays. E.C, F.P, P.B., and M.C., performed the RT-qPCR, and the WB. J.S performed the EU staining. C.B provided a valuable technical assistance. J.M.E provided valuable materials. F.C wrote the manuscript with input from I.D and E.C.

## MATERIAL AND METHODS

A full list of reagents including antibodies, commercial kits and oligonucleotides is supplied in Supplemental Table 1.

Access numbers for data generated in this paper are: GSE158118 and GSE158119

### Patients

Gene expressions in tumors and nevi were retrieved from several previously published datasets (including TCGA) indicated in the figure legends.

#### Cell culture and treatment

Cells were grown in 5% CO_2_ at 37°C in HAM-F10 (Gibco, Invitrogen) supplemented with 10% FCS and Penicillin-Streptomycin. Melanoma cell line 501mel was grown in 5% CO2 at 37°C in RPMI w/o HEPES (Gibco, Invitrogen) supplemented with 10% FCS and Gentamycin. Melanocyte cell line Hermes3A was grown in 10% CO_2_ at 37°C in RPMI w/o HEPES, supplemented with 10% FCS, Penicillin-Streptomycin, 200nM TPA (Sigma Aldrich), 200pM Cholera Toxin (Sigma Aldrich), 10ng/mL hSCF (Life Technologies), 10nM EDN-1 (Sigma Aldrich) and 2mM Glutamine (Invitrogen).

Cells were transfected with Lipofectamine RNAiMAX following the manufacture instructions with 25nM of siRNA ON-TARGETplus SMARTPool (Horizon Discovery) and cells were harvested 48 and/or 72h after transfection. All cell lines used were mycoplasm negative.

MMO99^MITF-SOX10-PAX3^ cells were generated as followed. Lentiviral vector pTET-SMP encoding human untagged MITF, SOX10 and PAX3 proteins was transduced in MM099 in presence of Polybrene and cells were selected with 3μg/ml of Puromycin. Conditional expression of pTET vector was carried out by adding 1μg/ml of Doxycycline in the medium for at least 24h.

#### CRISPR/Cas9 editing of 501mel^BIO-FLAG:CDK7^

501mel were co-transfected with vector px738 (encoding Cas9-HF-GFP and two guide RNAs targeting CDK7 locus and construct m599 (linear DNA fragment carrying homology regions to CDK7 locus and Puromycin-P2A-BIO-FLAG-CDK7N-termsequence (Key Resources Table) with transfection reagent FuGENE6. 24 h later single cells GFP positive were sorted in P96 well plates in presence of Puromycin (3μg/ml) with cell sorter. Single clones were let grown and selected for 4-6 weeks and surviving ones screened for gene editing through PCR using Phusion High-Fidelity TAQ Polymerase using different combination of primers (F1, F5, R3, R4, R5, see Key Resources Table). PCR positive clones were finally further amplified to perform western blot and Co-IPs validation.

#### Cell growth, cell death and senescence assay

To measure cell proliferation, cell death and senescence, cells were incubated first with CellTrace Violet according to the manufacture instructions. At the end of transfection and/or drug treatment, cells were incubated 1h with 100nM BafilomycinA1 and 2h with C12FDG; afterwards cells were rinsed and incubated 15 min with AnnexinV-APC. Cell proliferation, cell death and senescence were detected on a FORTESSA BD Biosciences Cytofluorometer. Data were analyzed with FlowJo software.

#### Reporter assay

The intGATA6r element was isolated by genomic PCR using Phusion High-Fidelity TAQ polymerase (ThermoFisher) with specific primers (see Key Resources Table). To allow the cloning within pCDNA-GFP vector, the first PCR product was further amplified by PCR with primers carrying MluI and SmaI restriction sites at the 5’ and 3’ respectively (MluI_F and SmaI_R primers in Key Resources Table). The immediate early CMV enhancer (ieEnh) in the pCDNA-GFP vector (pCDNA-ieEnh-CMV-GFP) was then replaced with the intGATA6r element (pCDNA-intGATA6r-CMV-GFP).

MITF-SOX10-PAX3 expression was induced in MMO99^MITF-SOX10-PAX3^ cells with doxycycline for 48h and cells were subsequently transfected with pCDNA-ieEnh-CMV-GFP or pCDNA-intGATA6r-CMV-GFP vectors for 24 h with FuGENE6 following manufacture instructions. RNA was then collected for qPCR and GFP protein signal was detected on cytofluorometer. FACS data were analyzed with FlowJo software.

#### Histology

Human tissue sections were de-paraffinized and dehydrated with Histosol and dilutions of ethanol (100%, 90%, 70% and 30%) and then rehydrated with demineralized water. Subsequently, sections were boiled in Sodium Citrate buffer (0.1M Citric acid, 0.1M Sodium citrate) for 15 min to unmask antigens. Alternatively, cells were growth on glass slides and fixed with 4% formaldehyde. Both tissues and cells were permeabilized with PBS and 0.1% TritoX-100. Blocking was done with 10% Fetal Bovin Serum before incubation with primary antibodies.

In situ hybridization of *ABCG2* mRNA was performed using the RNAscope assay according to manufacturer’s instructions (ACDBio). Cells and tissue sections were counterstained with DAPI and visualized using confocal microscope Spinning disk Leica CSU W1. Probes’ sequences were not provided by the manufacture.

#### EU incorporation assay

RNA labeling by EU incorporation was performed with Click-iT RNA Imaging kit following the manufacturer protocol. EU signal intensity was quantified using imaging system.

#### Cell survival assay

Normal or transfected cells were seeded at 5000 cells/well in a 96well plate and treated with increasing concentrations of THZ1, Vemurafenib or Trametinib. After 72h of incubation, cells were treated with PrestoBlue reagent according to manufacture instructions. The absorbance per well was measured with a microplate reader. The data were then analyzed using Prism8.

#### RT-qPCR

Total RNA was isolated from cells using a GenElute Mammalian Total RNA Miniprep kit (Sigma) and reverse transcribed with SuperScript IV reverse transcriptase (Invitrogen). The quantitative PCR was done using Lightcycler. The primer sequences for the different genes used in qPCR are indicated in Key Resources Table. The mRNA expression of the various analysed genes represents the ratio between values obtained from treated and untreated cells normalized with the housekeeping genes mRNA.

#### ChIP

Cells were grown on 15cm plates and, once reached 80% of confluence, were fixed with PBS + 0.4% formaldehyde solution for 10min. Fixation reaction was stopped with 2M Glycin pH 8. Cells were then pellet and suspended in lysis buffer (EDTA 10mM, Tris HCl pH8 50mM, SDS 1%) and sonicated with Covaris E220 AFA power 200Hz 6 cycles 200s to get a DNA fragmentation between 500 and 200bp. Chromatin was then diluted in 60ug aliquots with 8 volumes of ChIP dilution buffer (Tris HCl pH8 16.7mM, EDTA 1.2mM, NaCl 167mM, Triton X-100 1.1%, SDS 0.01%). The immuno-precipitations were done as following. 1-5ug of antibody was incubated overnight with chromatin and the complex antibody-chromatin was then captured with G protein sepharose beads (Sigma Aldrich) 1h at 4°C. Beads were washed 2 times with Low Salt Buffer (Trish HCl pH8 20mM,Ò EDTA 2mM, NaCl 150mM, Triton X-100 1%, SDS 0.1%), High Salt Buffer (Trish HCl pH8 20mM, EDTA 2mM, NaCl 500mM, Triton X-100 1%, SDS 0.1%), LiCl buffer (Tris HCl pH8 500mM, EDTA 1mM, Na deoxycholate 1%, NP40 1%, LiCl 0.25M), TE buffer (Tris HCl pH8 10mM, EDTA 1mM) and DNA was eluted 30min at room temperature with Elution buffer (NaHCO_3_ 0.1M, SDS 1%). DNA was finally purified through phenol-chloroform, re-suspended in 100uL of water and analysed by qPCR using a set of primers indicated in Key Resources Table.

#### RNA-seq

Rna-seq was performed like previously described (Laurette et al., 2019). For RNA-seq performed in this study, reads were preprocessed in order to remove adapter and low-quality sequences (Phred quality score below 20). After this preprocessing, reads shorter than 40 bases were discarded for further analysis. These preprocessing steps were performed using cutadapt version 1.10. Reads were mapped to rRNA sequences using bowtie version 2.2.8, and reads mapping to rRNA sequences were removed for further analysis. Reads were mapped onto the hg19 assembly of Homo sapiens genome using STAR version 2.5.3a. Gene expression quantification was performed from uniquely aligned reads using htseq-count version 0.6.1p1, with annotations from Ensembl version 75 and “union” mode. Only non-ambiguously assigned reads have been retained for further analyses. Read counts have been normalized across samples with the median-of-ratios method (Anders and Huber, 2010). Differential gene expression analysis was performed using the methodology implemented in the Bioconductor package DESeq2 version 1.16.1 (Love et al., 2014). P-values were adjusted for multiple testing by the method proposed by Benjamini and Hochberg (Benjamini and Hochberg, 1995). Deregulated genes were defined as genes with log2(foldchange) >1 or <-1 and adjusted p-value <0.05. Heatmaps were generated with the R package pheatmap v1.0.12.

#### ChIP-seq

Purified DNA fragments for ChIP-seq were prepared by using the ChIP-IT High Sensitivity Kit (Active Motif) and the related antibodies. ChIP-seq was performed on an Illumina sequencer as single-end 50 base reads following Illumina’s instructions. Image analysis and base calling were performed using RTA 1.17.20 and CASAVA 1.8.2. Reads were mapped onto the hg19 assembly of the human genome. Peak detection was performed using MACS (http://liulab.dfci.harvard.edu/MACS/) under settings where the input fraction was used as negative control. Peaks detected were annotated using HOMER (http://biowhat.ucsd.edu/homer/ngs/annotation.html) as well as TSS protein enrichment comparison. Quantitative comparison of RNA Pol II gene body enrichment was performed using seqMINER (http://bips.u-strasbg.fr/seqminer/). As reference coordinates, we used the MACS-determined peaks or the annotated TSS/TTS of human genes as defined by RefSeq database. Sequence enrichment was performed using RSAT (http://rsat.sb-roscoff.fr) with MACS-determined peaks as reference.

#### Bioinformatics

Expression matrix with row reads counts for the single cell experiment was retrieved from GEO (GSE116237). Then, data were normalized and clustered using the Seurat software package version 3.1.4 (Butler et al., 2018) in R version 3.6.1. Data were filtered and only genes detected in at least 3 cells and cells with at least 350 detected genes were kept for further analysis. Expression of 26,661 transcripts in 674 cells was quantified. To cluster cells, read counts were normalized using the method “LogNormalize” of the Seurat function NormalizeData. It divides gene expression counts by the total expression, multiplies this by a scale factor (10,000 was used), and log-transforms the result. Then, 2000 variable features were selected with the variance stabilizing transformation method using the Seurat function FindVariableGenes with default parameters. Integrated expression matrices were scaled (linear transformation) followed by principal component analysis (PCA) for linear dimensional reduction. The first 20 principal components (PCs) were used to cluster the cells with a resolution of 0.5 and as input to tSNE to visualize the dataset in two dimensions. The Bioconductor package AUCell v 1.6.1 (Aibar et al., 2017) was used to assess whether some cells from the Rambow dataset were enriched in gene sets of interest.

#### Gene Ontology

Gene ontology was performed using Metascape software developed by (Zhou et al., 2019).

## QUANTIFICATION AND STATISTICAL ANALYSIS

Statistical details of experimental can be found in figure legends or in the methods details

