## Supplemental informations for "TFIIH kinase CDK7 antagonizes phenotype switching and emergence of drug tolerance in melanoma"

**Supplemental data legends:**

**Supplemental Data 1:** List of the 261 genes up-regulated both in MM074$^{\text{CDK7i-R}}$ and MM047$^{\text{CDK7i-R}}$ cells (Page1). In this list, 35 genes are found in the GATA6 regulon (Page 2).

**Supplemental Data 2**: List of the 359 genes that composed the GATA regulon.

**Supplemental Data 3:** List of the 1767 genes that composed the SOX10 regulon**.**

**Supplemental Data 4:** Spearman's rank correlation coefficient between expression of *GATA6* and that of the indicated cluster markers extracted from TCGA bulk RNA-seq cohort of patient melanomas (skin cutaneous melanoma, n=444) (Pan-Cancer Atlas).

**Supplemental Table 1:** Table 1 provides a full list of reagents used in this manuscript including antibodies, commercial kits and oligonucleotides.

**Supplemental Figure 1: Generation of CDK7i- and BRAFi-resistant melanoma cells.**

**a.** Dose-response curves for the parental melanocytic-type MM074 and its CDK7i-resistant MM074$^{\text{CDK7i-R}}$ and BRAFi-resistant MM074$^{\text{BRAFi-R}}$ counterparts. Cells were treated with increasing doses of THZ1 for 72 h. Fractions of viable cells relative to DMSO-treated cells are shown. Error bars indicate mean values + SD for three biological triplicates. IC50 for each cell line is indicated.

**b.** Dose-response curves for parental melanocytic-type MM074, MM074$^{\text{BRAFi-R}}$ and MM074$^{\text{CDK7i-R}}$ cells. Cells were treated with increasing doses of Vemurafenib for 72 h. Fractions of viable cells relative to DMSO-treated cells are shown. Error bars indicate mean values + SD for three biological triplicates. IC50 for each cell line is indicated.

**c.** Parental MM047 cells and its CDK7i-resistant MM047$^{\text{CDK7i-R}}$ counterparts were treated as in panel **(a)**.

**d.** Dose-response curves for parental melanocytic-type MM074, MM074$^{\text{BRAFi-R}}$ and MM074$^{\text{CDK7i-R}}$ cells. Cells were treated with increasing doses of Trametinib for 72 h. Fractions of viable cells relative to DMSO-treated cells are shown. Error bars indicate mean values + SD for three biological triplicates. IC50 for each cell line is indicated.

**e-f.** Parental mesenchymal-like MM047 and its CDKi-resistant counterpart MM047$^{\text{CDK7i-R}}$ **(e)** or Parental melanocytic-type MM074 and its CDKi-resistant counterparts MM074$^{\text{CDK7i-R}}$ **(f)** were treated either with vehicle (DMSO) or THZ1 (50 nM) for 4 h. Transcribed RNAs were labelled by 5EU incorporation. The values of EU signal intensities are the means of three independent experiments + SD. N=2500 cells were analyzed in each experiment. P-value; ***<0.001, Student's t-test.

**Supplemental Figure 2: Phenotype switching of acquired CDK7i- and BRAFi-resistant melanoma cells**

**a.** GO was used to analyze the 261 genes up-regulated both in MM047$^{\text{CDK7i-R}}$ and MM074$^{\text{CDK7i-R}}$ cells. In the horizontal histograms the top 20 significant pathways obtained from the Metascape analysis are shown.

**b.** Boyden chamber assay for MM074 and MM074$^{\text{CDK7i-R}}$ cells. The graph represents the covered area + SD . P-value; ***<0.001, Student's t-test.

**c.** qRT-PCR analysis showing average *TBP*-normalized expression of *MLANA* in the indicated cell lines. Error bars indicate mean values + SD for three biological triplicates.

**d.** Image shows pellets of MM074, MM074$^{\text{CDK7i-R}}$ and MM074$^{\text{BRAFi-R}}$ cells. The pellets of MM074$^{\text{BRAFi-R}}$ cells have a brown color suggesting the presence of highly pigmented cells.

**Supplemental Figure 3: *GATA6* expression in melanoma *in vivo***

**a.** Venn diagram merging the up-regulated genes in MM074$^{CDK7i-R}$ and MM047$^{CDK7i-R}$ with a list of annotated transcription factors (TFs) (https://www.ncbi.nlm.nih.gov/pubmed/29425488).

**b.** The percentages of cells expressing each of the common 16 transcription factors identified above have been extracted from scRNA-seq data performed on 674 cells from a single drug-naïve PDX tumour (GSE116237) (Rambow et al., 2018).

**c.** Scatter plot expression values for *DMRTA1, MEIS3, GATA6, TLE4*, CASZ1, SETBP1 and *THRB* in nevi *vs* primary melanoma extracted from public bulk-RNAseq data of treatment-naïve melanocytic tumors (n=78) consisting of primary melanomas of the skin and benign melanocytic lesions (GSE98394) (Badal et al., 2017). P-values (Unpaired T-test) *<0.0332, **<0.0021, ***<0.0002 and ****<0.0001.

**Supplemental Figure 4: *ABCG2* is up-regulated in metastatic melanoma**

**a.** Venn diagram merging the up-regulated genes in MM074$^{CDK7i-R}$ and MM047$^{CDK7i-R}$ with the GATA6 regulon identified by pySCENIC.

**b**. Scatter plot expression values for *ABCG2* in primary *vs* metastatic melanoma extracted either from published DNA microarray data on a cohort of patient melanomas (skin cutaneous melanoma, n=83) (GSE8401) (Xu et al., 2008) (left panel) or from TCGA bulk RNA-seq data on a cohort of patient melanomas (skin cutaneous melanoma, n=444, Pan-Cancer Atlas) (right panel). P-values (Unpaired T-test) **<0.0021.

**c.** RNA FISH/mRNA *in situ* hybridization of *ABGC2*. **Top panel**: Probe was validated using 501mel and MM011 cells that do not express *ABCG2 vs.* MM029 that expresses it. **Bottom**

**panel**: Naevi and tumor sections (Primary melanoma or Cutaneous metastases) were used (n=3 sections from each tissue) with one representative micrograph from each tissue shown. Images were captured by confocal microscopy.

**d.** Venn diagram merging the up-regulated genes in MM074$^{\text{CDK7i-R}}$ and MM047$^{\text{CDK7i-R}}$ with a list of all ABC-transporters. Three ABC transporters are up-regulated in MM074$^{\text{CDK7i-R}}$ and/or MM047$^{\text{CDK7i-R}}$.

**e.** qRT-PCR analysis showing average *TBP*-normalized expression of *ABCB1, ABCC3* and *ABCG2* in the indicated cell lines. Error bars indicate mean values + SD for three biological triplicates.

**f.** qRT-PCR analysis showing average *TBP*-normalized expression of *ABCG2* in MM029 and MM099 melanoma cells treated with either siCTL or si*ABCG2*. Error bars indicate mean values + SD for three biological triplicates.

**Supplemental Figure 5: Characterization of 501mel$^{\text{BIO-FLAG:CDK7}}$ cells**

**a.** Protein lysates from 501mel and 501mel$^{\text{BIO-FLAG:CDK7}}$ cells were immuno-blotted for CDK7. Molecular sizes (KDa) of the proteins are indicated in kDa.

**b.** Immunoprecipitation was performed with anti-IgG, anti-CDK7 or anti-FLAG using whole cell extracts of 501mel and 501mel$^{\text{BIO-FLAG:CDK7}}$ cells. Proteins on the resin were resolved by SDS-PAGE and immunoblotted using anti-XPB or anti-XPD antibodies. Molecular sizes (KDa) of the proteins are indicated in kDa.

**Supplemental Figure 6: *SOX10* and *GATA6* expression level in individual cells treated with siSOX10**

Expression of *SOX10* and *GATA6* was determined in individual cell by AUCell from scRNA-seq performed on either the melanocytic-type MM074 **(a)** or MM087 **(b)** melanoma cells at different time points (0, 24, 48 and 72 h) post-transfection of si*SOX10* (GSE116237) (Rambow et al., 2018).

**Supplemental Figure 7: *GATA6* is over-expressed in metastatic melanoma**

Scatter plot expression values for *GATA6* in primary *vs* metastatic melanoma extracted either from published DNA microarray data on a cohort of patient melanomas (skin cutaneous melanoma, n=83) (GSE8401) (Xu et al., 2008) (left panel) or from TCGA bulk RNA-seq data on a cohort of patient melanomas (skin cutaneous melanoma, n=444, Pan-Cancer Atlas) (right panel). Unpaired T-test **<0.0021,.

**Supplemental Figure 8: *GATA6* is expressed in mesenchymal-like cells in tumors**

**a.** Seurat cluster Heatmap was generated from published scRNA-seq performed on a single drug-naïve PDX tumour (n=674 cells) (GSE116237) (Rambow et al., 2018). The Heatmap shows 9 different clusters into which the cells can be divided according to the expression of different referenced genes indicated in the left.

**b.** GO was used to analyze the genes characterizing each cluster identified above. The average P-value was retrieved for each cluster taking the 3 best GO per cluster, then z-score ((P-value of each biological process-average of P-value of each biological process)/standard deviation) was calculated. Clusters 7 and 8 represent mesenchymal-like and neural-crest cell-like melanoma cells, respectively.

**c.** The percentages cells expressing either mesenchymal marker *AXL* or the TF *GATA6* in each cluster identified above has been extracted from scRNA-seq performed on a single drug-naïve PDX tumour (n=674 cells) (GSE116237) (Rambow et al., 2018).

**Supplemental Table 1**

| REAGENT or RESOURCE | SOURCE | IDENTIFIER |
|---|---|---|
| **Antibodies** | | |
| Monoclonal Anti-Vinculin antibody produced in mouse | Sigma-Aldrich | V4505 |
| Polyclonal Anti-betaTubulin antibody produced in goat | Abcam | Ab21057 |
| Monoclonal Anti-SOX10 antibody produced in rabbit | Cell Signaling | D5V9L |
| Monoclonal Anti-MITF antibody produced in rabbit | Cell Signaling | D5G7V |
| Polyclonal Anti-PAX3 antibody produced in rabbit | ThermoFisher | 38-1801 |
| Monoclonal Anti-TFAP2A antibody produced in mouse | SC Biotechnology | sc-12726 |
| Monoclonal Anti-GATA6 antibody produced in rabbit | Cell Signaling | D61E4 |
| Monoclonal Anti-SOX9 antibody produced in rabbit | Cell Signaling | D8G8H |
| Monoclonal Anti-c-Jun antibody produced in rabbit | Cell Signaling | 9165 |
| Monoclonal Anti-RNAPII antibody produced in mouse | IGBMC, home made | 7C2 |
| Monoclonal Anti-pSer2 antibody produced in rat | Merck Millipore | 04-1571 |
| Monoclonal Anti-pSer5 antibody produced in rat | Merck Millipore | 04-1572 |
| Monoclonal Anti-pSer7 antibody produced in rat | Merck Millipore | 04-1570 |
| Polyclonal Anti-H3K27ac antibody produced in rabbit-ChIP Grade | Abcam | Ab4729 |
| Polyclonal Anti-HA antibody produced in rabbit-ChIP Grade | Abcam | Ab9110 |
| Monoclonal Anti-FLAG antibody produced in mouse | Merck Millipore | F3165 |
| **Biological Samples** | | |
| Sections of nevi and melanoma samples | Prof. B. Cribier, head of the *Laboratoire d'histopathologie et d'immunopathologie cutanées* at the Hôpital Civil of Strasbourg | N/A |

| | | |
|---|---|---|
| **Chemicals, Peptides, and Recombinant Proteins** | | |
| THZ1 | MedChemExpress | HY-80013 |
| Trametinib | MedChemExpress | HY-10999 |
| Vemurafenib | SelleckChem | PLX4032 |
| **Critical Commercial Assays** | | |
| Click-iT RNA Alexa Fluor 488 Imaging kit | Invitrogen | C10329 |
| PrestoBlue | ThermoFisher | A13262 |
| GenElute Mammalian Total RNA Miniprep kit | Merck | RTN70 |
| CellTrace Violet | ThermoFisher | C34571 |
| Phusion High-Fidelity TAQ polymerase | ThermoFisher | F-530XL |
| APC Annexin V | BD Biosciences | 550474 |
| BafilomycinA1 | Sigma-Aldrich | B1793 |
| FuGENE6 transfection reagent | Promega France | E2691 |
| Lipofectamine RNAiMAX reagent | ThermoFisher | 13778150 |
| **Experimental Models: Cell Lines** | | |
| Human; Short-term cultured melanoma cells | | (Gembarska et al., 2012) |
| Human; MM099$^{MITF-SOX10-PAX3}$ | This paper | N/A |
| Human; 501mel$^{BIO-FLAG:CDK7}$ | This paper | N/A |
| **Oligonucleotides** | | |
| Primer Actin, forward, ACATCTGCTGGAAGGTGGAC | This paper | N/A |
| Primer Actin, reverse, CCCAGCACAATGAAGATCAA | This paper | N/A |

| | | |
|---|---|---|
| Primer GAPDH, forward, ACAACTTTGGTATCGTGGAAGG | This paper | N/A |
| Primer GAPDH, reverse, GCCATCACGCCACAGTTTC | This paper | N/A |
| Primer MITF, forward, CATTGTTATGCTGGAAATGCTAGAA | This paper | N/A |
| Primer MITF, reverse, GGCTTGCTGTATGTGGTACTTGG | This paper | N/A |
| Primer TBP, forward, CGGCTGTTTAACTTCGCTTC | This paper | N/A |
| Primer TBP, reverse, CACACGCCAAGAAACAGTGA | This paper | N/A |
| Primer SOX10, forward, CCAGTTTGACTACTCTGACCATCAG | This paper | N/A |
| Primer SOX10, reverse, ATATAGGAGAAGGCCGAGTAGAGG | This paper | N/A |
| Primer ABCC3, forward, GGAAAACGTGCTTTTCGGCAA | This paper | N/A |
| Primer ABCC3, reverse, CCCCCAGACAGGTTAATGCC | This paper | N/A |
| Primer ABCB1, forward, GGAGGCCAACATACATGCCT | This paper | N/A |
| Primer ABCB1, reverse, AGGCTGTCTAACAAGGGCAC | This paper | N/A |
| Primer ABCG2, forward, TCAGGAGGCCTTGGGATACT | This paper | N/A |
| Primer ABCG2, reverse, GTCTTCTTCTCTGTTTAATGCCACA | This paper | N/A |
| Primer GATA6, forward, ACCACCTTATGGCGCAGAAA | This paper | N/A |
| Primer GATA6, reverse, ATAGCAAGTGGTCTGGGCAC | This paper | N/A |
| Primer VIPR1, forward, GCAGACCATGTTCTACGGTTC | This paper | N/A |
| Primer VIPR1, reverse, AACAGGCTCAGGATAGCTGTG | This paper | N/A |
| Primer ERCC2, forward, GGCGCCATGAAGCTCAAC | This paper | N/A |
| Primer ERCC2, reverse, TCCAGGACTCCATGACCCTT | This paper | N/A |
| Primer for ChIP, GATA6, forward, GAAAAAGCCGTAAGCACAGTCTCA | This paper | N/A |
| Primer for ChIP, GATA6, reverse, ACGACTGTGTGATCCTTCCCA | This paper | N/A |
| Primer for ChIP, TYR, forward, CATCCTTCTGTAAGGCCACAG | This paper | N/A |

| | | |
|---|---|---|
| Primer for ChIP, TYR, reverse, ACTGGGAATGAAGGGCAAG | This paper | N/A |
| Primer for ChIP, PRMG, forward, ACAGAGCGACACCCTGTCAT | This paper | N/A |
| Primer for ChIP, PRMG, reverse, AGGCGGTGGTTACACAACA | This paper | N/A |
| gRNA1, CDK7, tcgggctttacggcgccgga | This paper | N/A |
| gRNA2, CDK7, acttcacgtccagagccatc | This paper | N/A |
| Puromycin-P2A-BIO-FLAG-CDK7N-termsequence:ctttaaattcgtgttgtcctgggagctcgcccttttcggctggagtcgggctttacggcgCCGgATGACCGAGTACAAGCCCACGgtgcgcctcgccacccgcgacgacgtccccagggccgtacgcaccctcgccgccgcgttcgccgactaccccgccacgcgccacaccgtcgatccggaccgccacatcgagcgggtcaccgagctgcaagaactcttcctcacgcgcgtcgggctcgacatcggcaaggtgtgggtcgcggacgacggcgccgcggtggcggtctggaccacgccggagagcgtcgaagcgggggcggtgttcgccgagatcggcccgcgcatggccgagttgagcggttcccggctggccgcgcagcaacagatggaaggcctcctggcgccgcaccggcccaaggagcccgcgtggttcctggccaccgtcggcgtctcgcccgaccaccagggcaagggtctgggcagcgccgtcgtgctccccggagtggaggcggccgagcgcgccggggtgcccgccttcctggaAacctccgcgccccggaacctccccttctacgagcggctcggcttcaccgtcaccgccgacgtcgaggtgcccgaaggaccgcgcacctggtgcatgacccgcaagcccggtgccGGaagcggagctactaacttcagcctgctgaagcaggctggagacgtggaggagaaccctggacctggcctgaatgacatctttgaggcccagaagatcgagtggcatgagggaggaatggactacaaggacgacgatgacaagggaggcggagggagtggaggcggtggCAGCGGTGGCGGAGGGAGTGctctggacgtgaagtctcgggcaaagcgttatgagaagctggacttccttggggagggacaggtg) | This paper | N/A |
| Single clones check, F1: GAACGCCAACCGCCTGG | This paper | N/A |
| Single clones check, F5: AAGAACTCTTCCTCACGCGCG | This paper | N/A |
| Single clones check, R3: CCGAGACTTCACGTCCAGAGC | This paper | N/A |
| Single clones check, R4: AAACGTGGCGGGTCAGTCTCC | This paper | N/A |
| Single clones check, R5: CCTTCCATCTGTTGCTGCGC | This paper | N/A |
| MluI_F:aaaaACGCGTACGATTTGCAGGAAAGCATTTCCC | | |
| SmaI_R:aaaaCCCGGGCATGGGTT CTTCGGCTTGTGG | | |
| intGATA6r forward primer ACGATTT GCAGGAAAGCATTTCCC | | |

| | | |
|---|---|---|
| intGATA6r reverse primer CATGGGTTCTTCGGCTTGTGG | | |
| **Recombinant DNA** | | |
| pCDNA-ieEnh-CMV-GFP | This paper | N/A |
| pCDNA-intGATA6r-CMV-GFP | This paper | N/A |
| pTET-SMP-MITF/SOX10/PAX3 | This paper | N/A |
| pCDNA-GFP | Thermofisher | |
| **Software and Algorithms** | | |
| Metascape software | (Zhou et al., 2019) | |
| AUCell v 1.6.1 | (Aibar et al., 2017) | |
| Seurat software package version 3.1.4 | (Butler et al., 2018) | |
| RSAT | (http://rsat.sb-roscoff.fr) | |
| seqMINER | (http://bips.u-strasbg.fr/seqminer/) | |
| HOMER | (http://biowhat.ucsd.edu/homer/ngs/annotation.html) | |
| MACS | (http://liulab.dfci.harvard.edu/MACS/) | |
| **Other** | | |
| FACSAria™ Fusion cell sorter | BD Biosciences | |
| LSRFortessa™ Flow Cytometer | BD Biosciences | |
| INCell Analyzer 1000 imaging system | GE Healthcare | |
| Mithras LB 940 microplate reader | Berthold | |
| LightCycler® 480 Instrument II | Roche Diagnostic | |

| | |
|---|---|
| HiSeq2500 system | Illumina |

### Supplemental Figure 1

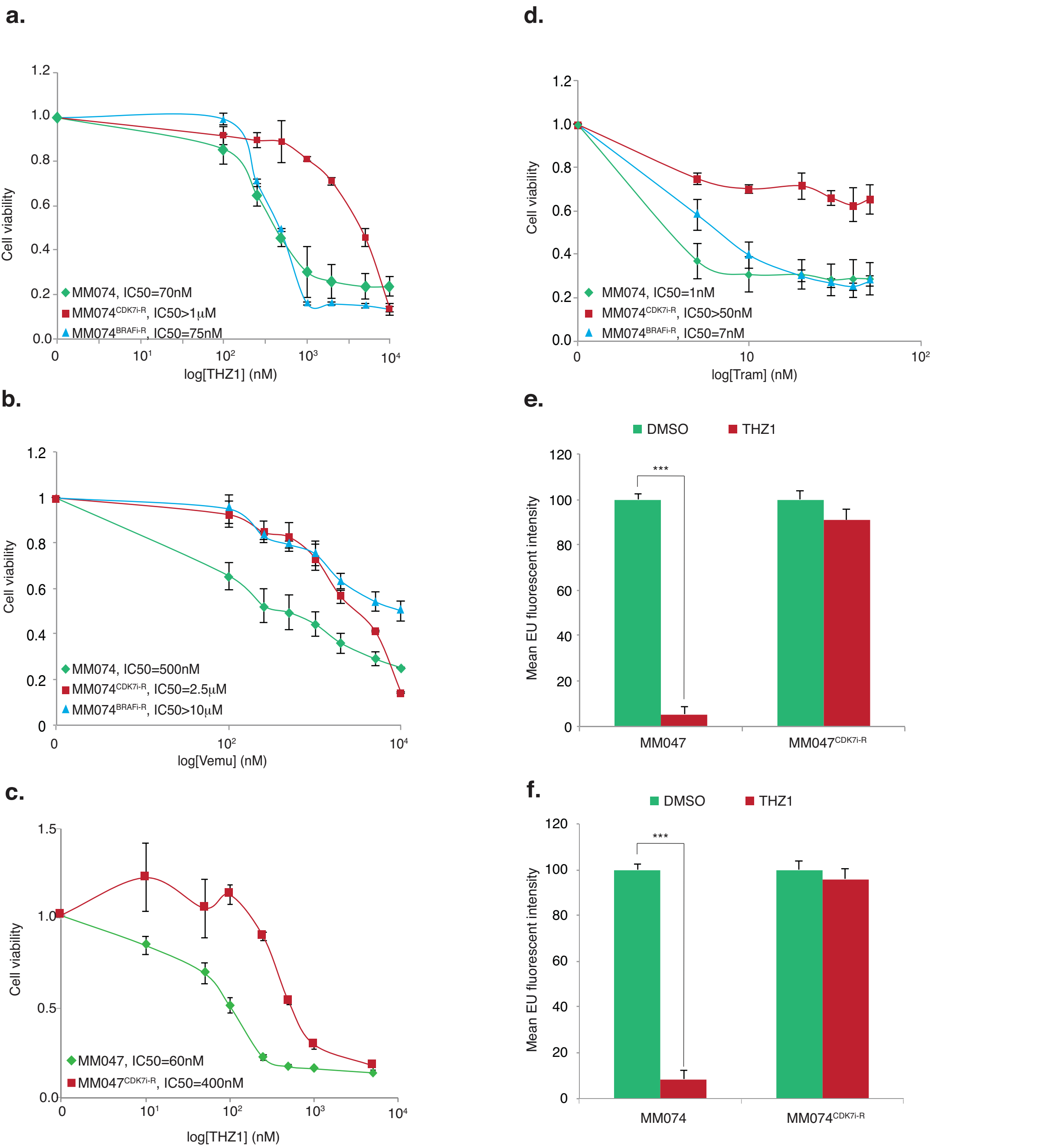

### Supplemental Figure 2

a.

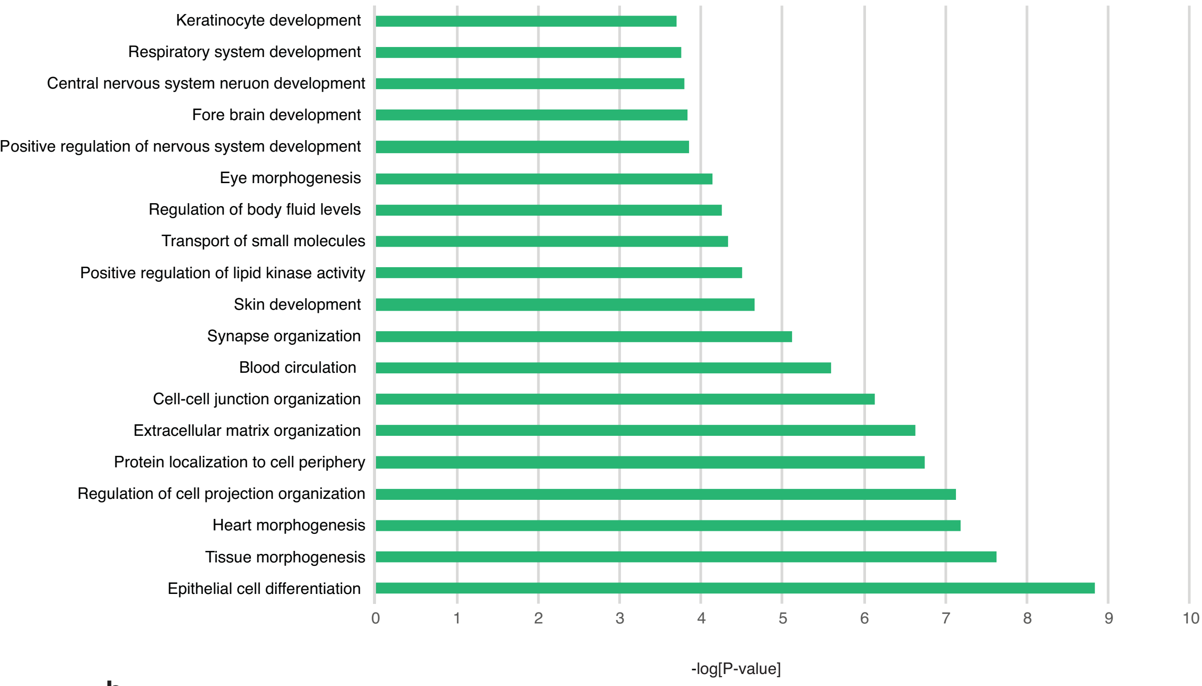

b.

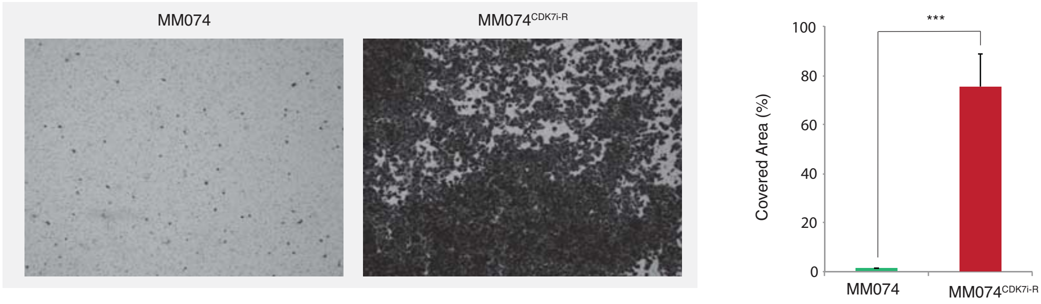

c.

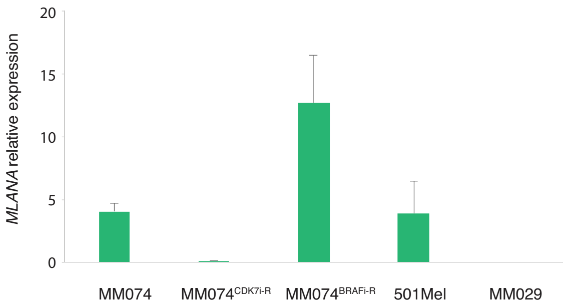

d.

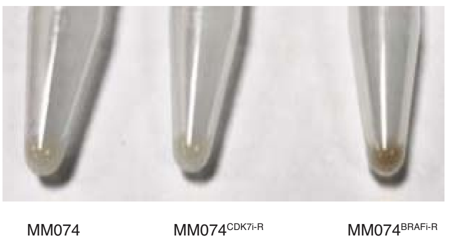

### Supplemental Figure 3

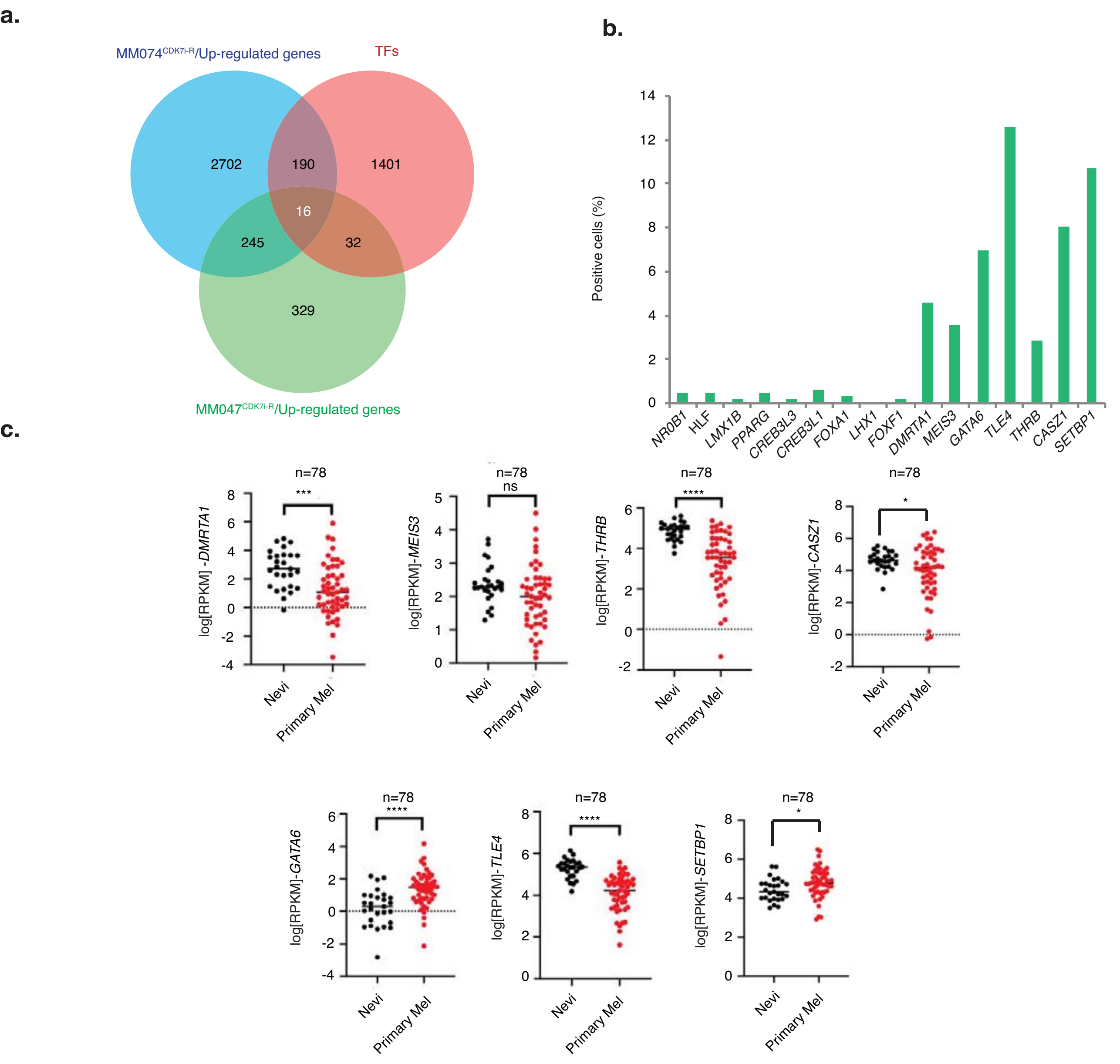

#### Supplemental Figure 4

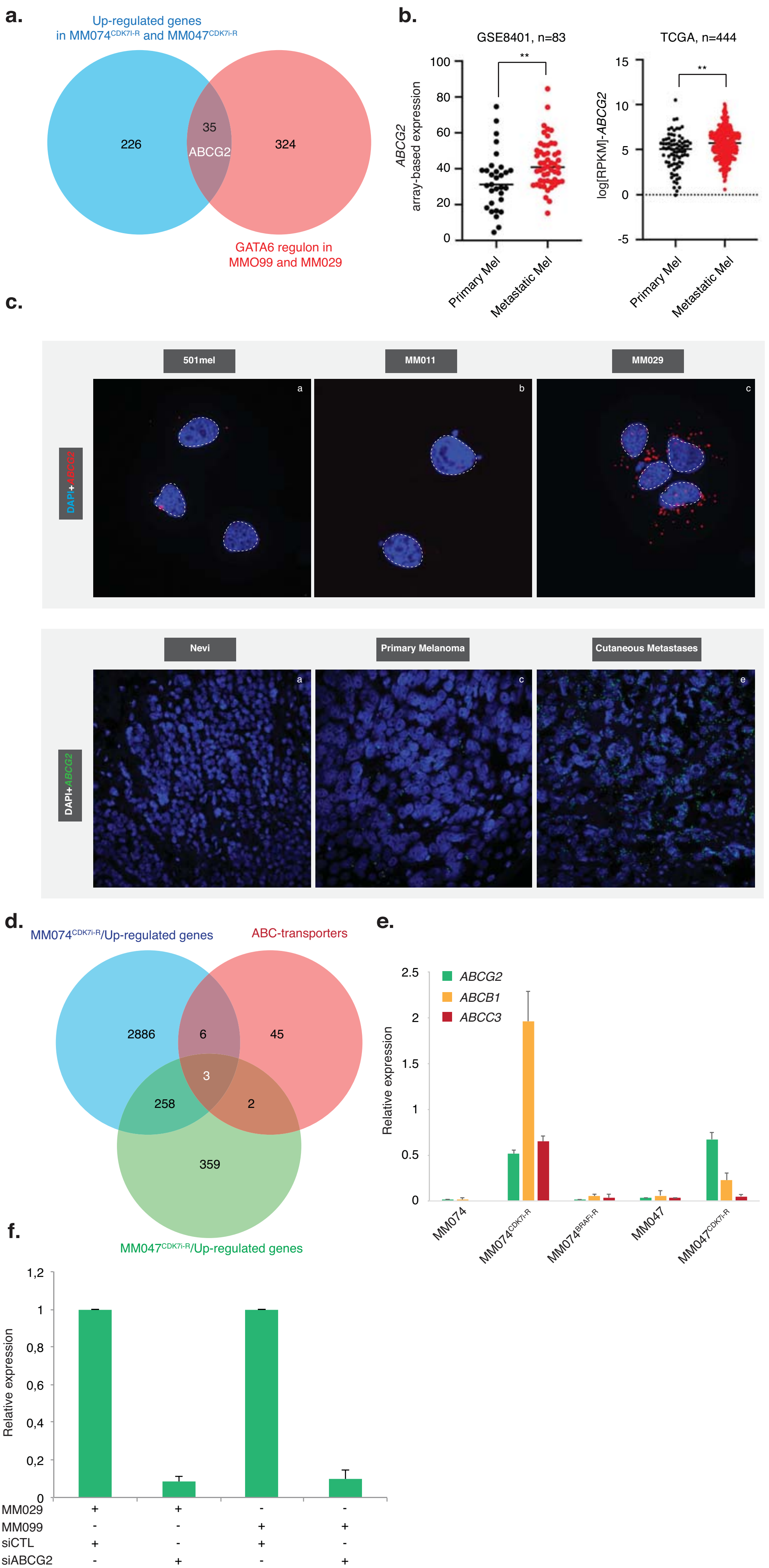

### Supplemental Figure 5

**a.**

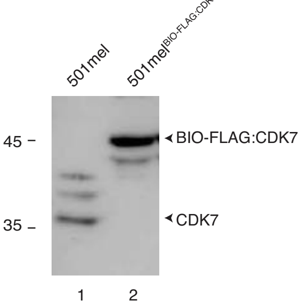

**b.**

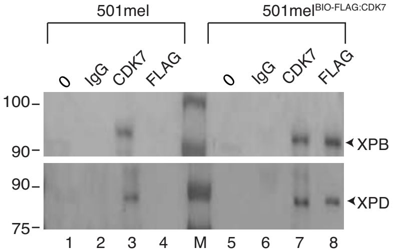

### Supplemental Figure 6

**a.**

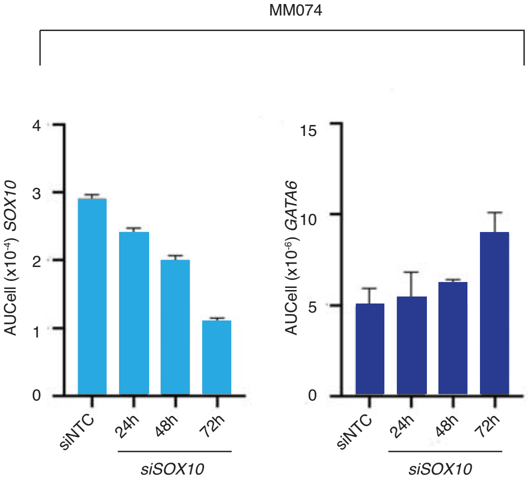

**b.**

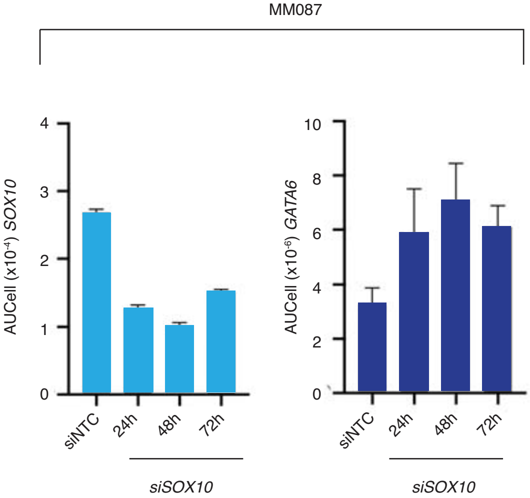

### Supplemental Figure 7

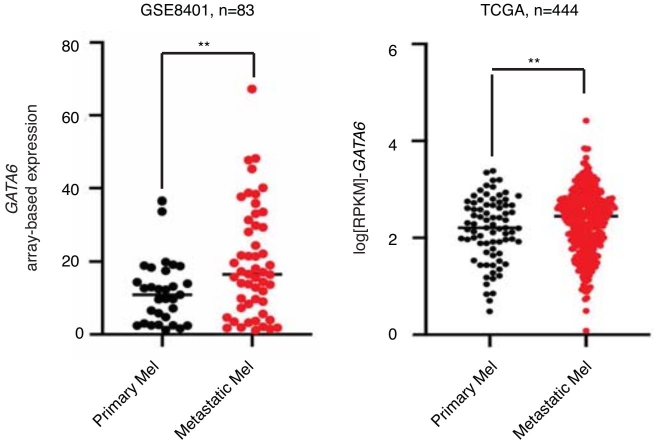

### Supplemental Figure 8

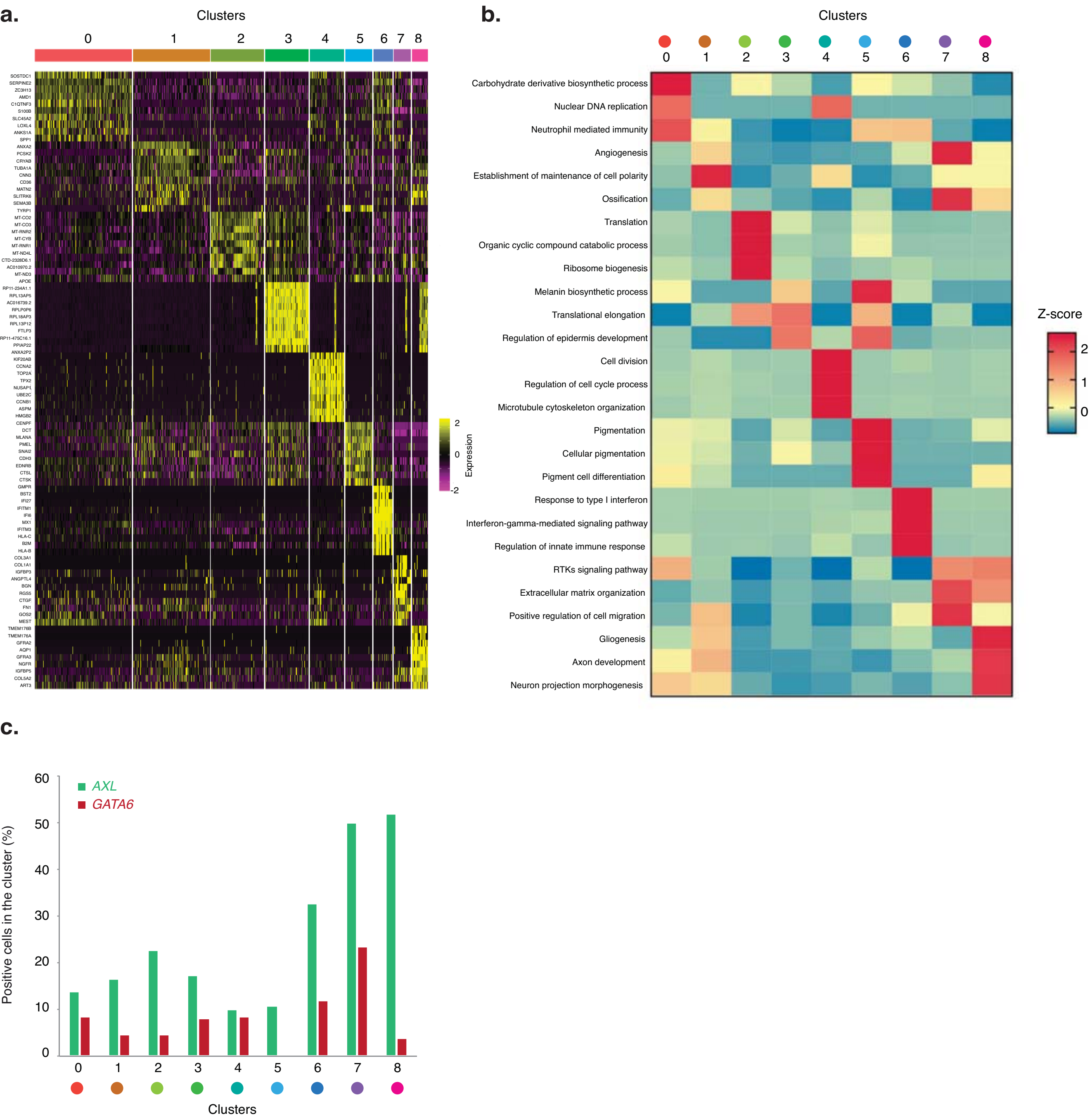
